# FLEXIQuant-LF: Robust Regression to Quantify Protein Modification Extent in Label-Free Proteomics Data

**DOI:** 10.1101/2020.05.11.088492

**Authors:** Konstantin Kahnert, Christoph N. Schlaffner, Jan Muntel, Ruchi Chauhan, Bernhard Y. Renard, Judith A. Steen, Hanno Steen

## Abstract

Improvements in LC-MS/MS methods and technology have enabled the identification of thousands of modified peptides in a single experiment. However, protein regulation by post-translational modifications (PTMs) is not binary, making methods to quantify the modification extent crucial to understanding the role of PTMs. Here, we introduce FLEXIQuant-LF, a software tool for large-scale identification of differentially modified peptides and quantification of their modification extent without prior knowledge of the type of modification. We developed FLEXIQuant-LF using label-free quantification of unmodified peptides and robust linear regression to quantify the modification extent of peptides. As proof of concept, we applied FLEXIQuant-LF to data-independent-acquisition (DIA) data of the anaphase promoting complex/cyclosome (APC/C) during mitosis. The unbiased FLEXIQuant-LF approach to assess the modification extent in quantitative proteomics data provides a better understanding of the function and regulation of PTMs. The software is available at https://github.com/SteenOmicsLab/FLEXIQuantLF.

## Introduction

Most cellular processes are regulated by post-translational modifications (PTMs). Given the sensitivity of current mass spectrometric analyses, even non-functional basal PTMs can be identified by mass spectrometry, thus an understanding of the stoichiometry is crucial to understanding function. Numerous bioinformatics tools that enable unbiased and large-scale identification of modified peptides and localization of the modification site such as MSFragger (1), MetaMorpheus (2) or TagGraph (3) have been developed. With steadily increasing qualitative information about the identity of PTMs (4) (5) (6) (7) (8) (9) (10), tools to investigate the modification quantity, i.e. the proportion of the protein carrying PTMs, become increasingly important. Thus far, analysis of the PTM extent has mainly focused on quantifying the modified peptides, e.g. using PTM enrichment methods (11) (12) (13), heavy isotope labeled synthetic peptides (14) or by enzymatically removing (15) (16) PTMs. These approaches require prior knowledge of the PTMs and have been primarily applied to the phosphoproteome.

The development of FLEXIQuant (*F*ull-*L*ength-*Ex*pressed Stable *I*sotope-labeled Proteins for *Quant*ification) (17) (18), for the first time, enabled the quantification of the extent of modification across a whole protein without prior knowledge of modification identities. FLEXIQuant relies on the principle that the total number of molecules of a given protein is conserved in a sample, thus the number of molecules of each unique peptide derived from that protein will be equal. If a peptide is modified chemically by a PTM this would reduce the abundance of its unmodified cognate in the peptide pool. In FLEXIQuant, this idea is realized by analyzing the peptide pool of the sample with an added unmodified heavy isotope-labeled full-length reference protein. The extent of modification is calculated by comparing the quantities of labeled reference peptides derived from the standard to the unlabeled cognate peptides from the sample, i.e. the unlabeled endogenous protein. The advantage of this approach is the precise and accurate quantification of the degree of modification of each quantified peptide and the ability to calculate the absolute concentration. FLEXIQuant and derivatives thereof have, for instance, been applied to study Tiki1 modification in head formation in frogs (19), to gain insights into the phosphorylation dynamics of GSK3β-dependent phosphorylation of DCX (20), the cell cycle dependent phosphorylation of KifC1 (21) or to investigate post-translational modification of Tau in Alzheimer’s disease (22). However, the limitation of this method is the requirement of a purified, unmodified, labeled reference protein and thus, it is labor intensive and has largely been used to investigate individual proteins. Recently, a PTM analysis pipeline without the need for an internal reference protein has been published (23). While this is encouraging, the quantification relies on isobaric labeling approaches which are expensive to conduct due to the high cost of the labeling reagents and is therefore not widely applicable, especially not for large-scale experiments. This highlights the need for bioinformatic tools suitable for the analysis of label-free experiments.

Here, we introduce FLEXIQuant-LF as an unbiased, label-free computational tool to detect modified peptides and to quantify the degree of modification based solely on the unmodified peptide species building upon the FLEXIQuant (17) idea developed in our lab. We developed this approach to identify and elucidate differential protein modification in commonly analyzed time series or case-control studies. In these studies, a timepoint or control group is used as reference to enable a FLEXIQuant-like modification analysis. The key requirement of FLEXIQuant-LF is the unbiased, comprehensive and highly reproducible quantification of peptides, therefore, we use a data-independent acquisition (DIA) strategy (namely, SWATH (24)). Here, we demonstrate the utility of FLEXIQuant-LF by applying our method to interrogate the modification dynamics of the anaphase-promoting complex/cyclosome (APC/C) during mitosis (17) (25).

## Materials and Methods

### FLEXIQuant-LF algorithm and benchmarking

FLEXIQuant-LF identifies modified peptides, i.e. peptides whose intensities strongly deviate from those of a reference sample, using robust linear regression. Based on the distance of each peptide to the regression line, Relative Modification (RM) scores are calculated that, in analogy to the light to heavy ratio of the original FLEXIQuant method, correspond to the degree of modification (**Fig. 1B**).

**Figure 1:**
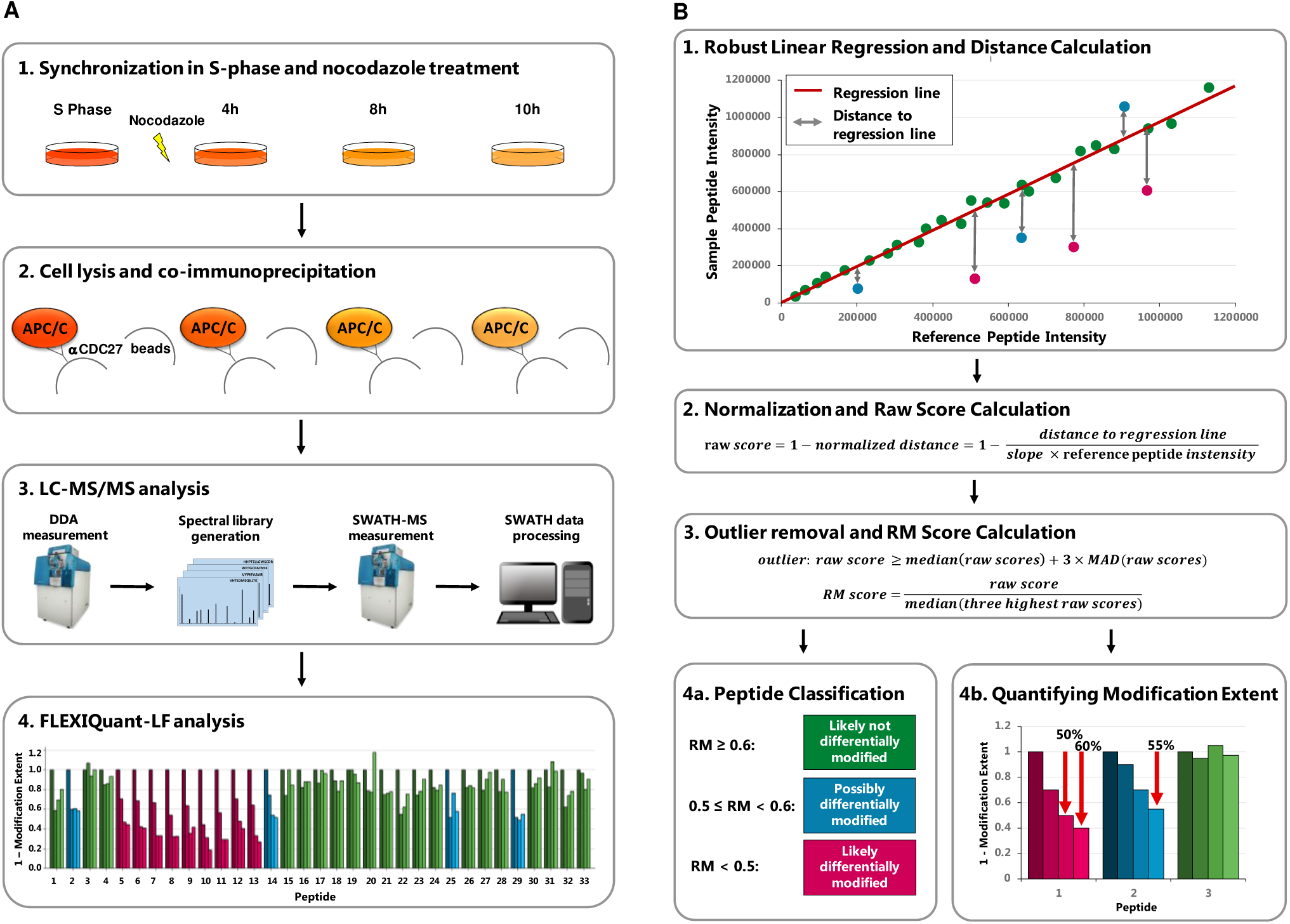
Workflow and FLEXIQuant-LF Concept. (A) Workflow. A1) HeLa cells were synchronized in S-phase using thymidine (dark orange). Upon release from thymidine block, cells were treated with nocodazole and samples were collected after 4h (medium dark orange), 8h (orange) and 10h (light orange). A2) APC/C was co-immunoprecipitated using an anti-CDC27 antibody. A3) All samples were trypsinized separately and analyzed by LC-MS/MS in DDA mode to generate a spectral library, which was subsequently analyzed by SWATH-MS (see **Supplementary Table 1** for raw peptide intensities of all quantified APC/C proteins). A4) FLEXIQuant-LF-based differential modification analysis of APC/C proteins (see **Supplementary Table 2** for all resulting RM scores). (B) FLEXIQuant-LF overview. B1) Firstly, a RANSAC-based robust linear regression model is fitted to the intensities of unmodified peptide species using a reference sample as independent variable and the sample of interest as dependent variable and the vertical distance to the regression line of each peptide is determined. B2) For each peptide, the distance is then normalized by dividing by the slope of the regression line multiplied by the intensity of the peptide in the reference sample and the result is subtracted from 1 to yield raw scores. B3) Peptides with a raw score above three standard deviations of the median (MAD) of all raw scores are classified as outliers and excluded from the subsequent RM score calculation. The remaining raw scores are then scaled using the median of the three highest raw scores resulting in a metric, termed RM score which is equal to 1 minus the extent of modification. B4) Lastly, peptides are classified in three categories based on their RM scores and the extent of modification is visualized: i) RM score < 0.5: peptide is likely differentially modified (magenta bars), ii) 0.5 ≤ RM score < 0.6 : peptide is possibly differentially modified (blue bars) and iii) RM score ≥ 0.6: peptide is likely not differentially modified (green bars).

More specifically, the FLEXIQuant-LF algorithm trains a linear regression model for each sample in the input file using the random sample consensus (RANSAC) algorithm (26) to identify outliers iteratively and fit the model only based on the inliers, i.e. based on unmodified peptides. For this, peptide intensities of each sample are used as dependent variable (*y*_*i*_), peptide intensities of a user defined reference sample as independent variable (*x*_*i*_) and no intercept is fitted (**Fig. 1 B1**):

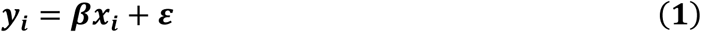

In Formula 1, the slope of the regression line is described by ***β*** whereas ***ε*** represents an error term. To increase reproducibility and enhance the fit of the linear regression model to the data the algorithm is executed 30 times and the best model, based on *r*^2^ scores, is selected (more detailed information in **Supplementary Reproducibility**). Subsequently, vertical distances between each peptide intensity and the obtained regression line are determined, i.e. intensity differences between measured and expected peptide intensities. To correct for sample-specific differences in individual protein abundance and peptide-specific differences in intensity, distances are normalized by dividing them by the expected intensity, i.e. the reference intensity of the peptide from the independent variable multiplied by the slope of the regression line. FLEXIQuant-LF raw scores are then calculated by subtracting the normalized distances from 1 (**Fig. 1 B2**). For each sample, peptides with a raw score above three standard deviations of the median (MAD) of all raw scores are classified as outliers and excluded from the subsequent RM score (Relative modification score) calculation. RM scores are calculated by dividing each inlier raw score by the median of the three highest raw scores (after removing outliers) for each sample, corresponding to one minus the extent of modification as defined by FLEXIQuant (**Fig. 1 B3**, detailed description in **Supplementary Methods**).

The FLEXIQuant-LF method was implemented in Python 3.7.3 using Scikit-learn 0.21.2 (27), NumPy 1.16.4 (28), Pandas 0.24.2 (29), SciPy 1.2.1 (30), Matplotlib 3.1.0 (31), Seaborn 0.9.0 (32). Parameters of the RANSAC algorithm are set as FLEXIQuant-LF defaults as follows: max_trials=1000, base_estimator=linear_model.LinearRegression, min_samples=0.5, stop_probability=1, loss=“squared_loss”, residual_threshold=MAD(sample intensities)^2^. The Fit_intercept parameter of the linear regression algorithm is set to False. The graphical user interface was built using QT Designer 5.13.0 (33), QT 5.9.7 (34) and PyQT 5.13.2 (35). PyInstaller 3.6 (36) was used to create the executable file. The command line interface version was built utilizing Click 7.0 (37).

To benchmark FLEXIQuant-LF and identify differentially modified peptides in the time course experiment, peptide intensities of the proteins APC1, APC4, APC5, APC7, APC10, APC15, APC16, CDC16, CDC20, CDC23, CDC26 and CDC27 from the well-studied APC/C were extracted. Peptides used for quantification had to be unmodified, or only modified with oxidation on methionine and/or carbamidomethylation on cysteine residues. Furthermore, they had to be quantifiable in all LC/MS runs. Median intensities of all replicates of each time point were calculated. Time point 0 (S phase) was defined as reference time point for FLEXIQuant-LF application using the default values described above. Peptides were then classified in three categories based on their RM scores at time point 10h: **1)** RM score < 0.5: peptide is likely differentially modified (magenta bars), **2)** ≤ RM score < 0.6: peptide is possibly differentially modified (blue bars) and **3)** RM score ≥ 0.6: peptide is likely <u>not</u> differentially modified (green bars) (**Fig. 1 B4**).

### Application improvement using a “superprotein” approach

FLEXIQuant-LF results improve the more peptides are available to determine the linear regression. Two proteins (APC15, APC16) of APC/C are small and thus resulted in fewer than five quantified peptides in our experiments. To extend our analysis of APC/C to these proteins with an insufficient number of quantified peptides, we approached APC/C as a “superprotein” drawing on the well-studied equal stoichiometry of APC/C components in the complex (with the exception of CDC20, which was omitted from this “superprotein” approach, as it is known to change in abundance during M-phase (see results)). Therefore, all reliably quantified peptides were analyzed together as if there were derived from a pan-APC/C “superprotein” following the steps described above for single proteins up to raw score calculation. Peptides were then split according to the proteins they originate from and RM scores were subsequently calculated for each protein separately to improve quantification accuracy.

### Sample generation and co-IP

HeLa S3 cells (ATCC, Manassas, VA) were grown in DMEM media (Invitrogen, Carlsbad, CA) supplemented with 10% FBS (Invitrogen), 2 mM l-glutamine (Invitrogen), 100 μg/ml penicillin and streptomycin mix (Invitrogen). For S phase, cells were treated with 2 mM thymidine (Sigma, St. Louis, MO) at 80% confluency for 20 hours and cultured for 8 hours in media without thymidine. For M phase, thymidine-treated cells were cultured for 3 hours in fresh media and treated with 100 ng/ml nocodazole (Sigma) and were collected at 4 h, 8 h and 10 h time points. Cells were washed with PBS and lysed in 1x RIPA lysis buffer (Santa Cruz, Dallas, TX) supplemented with 1 M DTT (Sigma), 10% glycerol, protease and phosphatase inhibitors (HALT, Thermo Fisher Scientific, Waltham, MA) using a bead beater homogenizer (Precellys, Rockville, MD). Cell lysate volumes of 300 µl with 2-3 mg protein were precleared with 20 μl Affi-Prep Protein A beads (Biorad, Hercules, CA) and 20 μg mouse IgG (Santa Cruz) for 2 h at 4°C with end-on-end rotation. Affi-Prep Protein A beads were washed twice with 6 volumes of buffer A (10 mM phosphate buffer, pH 7.4), 120 mM NaCl, 2.7 mM KCl, 0.1% Triton-X), twice with 6 volumes of buffer B (10 mM phosphate buffer, pH 7.4, 120 mM NaCl, 300 mM KCl, 0.1% Triton-X) and then once more with 6 volumes of buffer A. Freshly prepared beads (20 μl) conjugated with 20 μg CDC27 antibody (AF3.1, Santa Cruz) were incubated with precleared lysates for 2 h at 4°C with end-over-end rotation. As control, the experiment was repeated using beads without conjugated antibody (controls). The beads were washed as described above. The immunoprecipitate was eluted from beads by boiling in 1x Laemmli buffer with β-mercaptoethanol.

### Sample preparation/digestion

The samples were separated on a 4-12% SDS-PAGE gel (Invitrogen) in 1x MES buffer (Invitrogen) and the excised band was subjected to in-gel digestion. In brief, proteins were reduced with DTT, alkylated with iodoacetamide and digested with trypsin. After extraction of the peptides with acetonitrile, peptides were dried down and resuspended in a buffer containing 5% formic acid/5% acetonitrile/90% water. HRM calibration peptides (Biognosys, Schlieren, Switzerland) were added to the samples prior to analysis according to manufacturer instructions.

### Spectral library generation

To generate a spectral library, we analyzed each sample once using a standard data-dependent acquisition (DDA) method on a TripleTOF 5600 mass spectrometer (Sciex, Framingham, MA) coupled online to a nanoLC system (Sciex/Eksigent, Dublin, CA), equipped with an LC-chip system (cHiPLC nanoflex, Eksigent, trapping column: Nano cHiPLC trap column 200 μm × 0.5 mm Reprosil C18 3 μm 120 Å, analytical column: Nano cHiPLC column 75 μm × 15 cm Reprosil C18 3 μm 120 Å). Peptides were separated by a linear gradient from 95% buffer A (0.2% FA in water) / 5% buffer B (0.2% FA in ACN) to 70% buffer A / 30% buffer B within 90 min. The mass spectrometer was operated in data-dependent TOP50 mode with the following settings: MS1 mass range 300-1,700 Th with 250 ms accumulation time; MS2 mass range 100-1,700 Th with 50 ms accumulation time and following MS2 selection criteria: UNIT resolution, intensity threshold 100 cts; charge states 2-5. Dynamic exclusion was set to 17 s.

The DDA data were searched with MaxQuant (v1.5.2.8) (38) against the human UNIPROT database (only reviewed entries, downloaded on Oct. 31^st^, 2014) using the .WIFF files without additional file conversion. The protein sequence database was appended with common laboratory contaminants (cRAP, version 2012.01.01) and the iRT fusion protein sequence (39) (Biognosys) resulting in 20,296 entries. The following settings were applied: trypsin with up to 2 missed cleavages; mass tolerances set to 0.1 Da for the first search and 0.01 for the main search. Oxidation of M (+15.995 Da), phosphorylation of S, T, Y (+79.966 Da) and acetylation of N-TERM (+42.011 Da) were selected as dynamic modifications and carbamidomethylation of C (+57.021 Da) was selected as a static modification. FDR was set to 1% at both the peptide and protein level. The default settings were used for all other search parameters. Finally, a spectral library based on the MaxQuant search results was generated in Spectronaut 7.0 (Biognosys) using the following settings: Q value cut-off of 0.01, and a minimum of 3 and a maximum of 6 fragment ions.

To improve PTM identification output, the DDA data were also searched in Protein Pilot (v4.5.1, Sciex) using the same database described above and the following settings: sample type – identification, cys alkylation – iodoacetamide, digestion – trypsin, instrument – TripleTOF 5600, special factors – phosphorylation emphasis and gel-based ID, ID focus – biological modifications, search effort – thorough ID. For the ProteinPilot searches, it is not necessary to select the instrument mass accuracy, number of missed cleavages or the PTMs. The search results were filtered by a peptide FDR of 1%.

### SWATH sample acquisition

The SWATH data were acquired using the same mass spectrometer (TripleTOF 5600, Sciex) and LC-setup as described for the DDA samples. The mass spectrometer was operated in SWATH mode covering the mass range from 400 to 1,000 Th with 75 windows (8 Th width with 1 Da overlap). The accumulation time was set to 40 ms. Additionally, an MS1 scan was acquired in the mass range from 400-1,000 Th with an accumulation time of 250 ms, resulting in a cycle time of 3.3 s. The samples were acquired in triplicates.

### SWATH data analysis

All SWATH data were directly analyzed in Spectronaut 7.0 (Biognosys) (40) without any file conversion using the previously generated spectral library. The following settings were applied in Spectronaut 7.0: peak detection – dynamic iRT, correction factor 1; dynamic score refinement and MS1 scoring – enabled; interference correction and cross run normalization (total peak area) – enabled. Peptides were grouped according to the protein grouping by MaxQuant during generation of the spectral library. Spiked-in HRM peptides were used in the analysis for retention time and m/z calibration. The m/z tolerance was in the range of 12 ppm and the median extraction window was 4 min. All results were filtered by a Q value of 0.05 (equivalent to an FDR of 5% at the peptide level). All other settings were set to default. Protein intensities were calculated by summing the peptide peak areas (sum of fragment ion peak areas as calculated by Spectronaut) from the Spectronaut output file.

## Results

The objective of our study was to establish a workflow to elucidate modification dynamics in an unbiased manner without the need for heavy isotope-labeled reference proteins or peptides. Our strategy focused on profiling unmodified peptides, as any variation in the extent of modification of a peptide results in the reduction of the amount/intensity of the remaining unmodified peptide species. This consideration also applies to modifications on lysine or arginine residues or close to tryptic cleavage sites that interfere with the tryptic cleavage. In such cases, the affected tryptic cleavage yield shows a reduction which is equivalent to the modification extent. This interference with the trypsin cleavage results in a proteolytic missed cleaved peptide, which in turn leads to a change in the amount/intensity of the two unmodified, fully cleaved peptide species. Thus, variation in the amount of an unmodified peptide can be used to identify and quantify peptides whose modification state changes relative to a reference point in a time series or in a dose-dependent manner. The idea of focusing on the quantity of the unmodified peptide to identify and quantify modified peptides was implemented in the original FLEXIQuant approach, which uses stable isotope-labeled reference samples (17) (18). Here, we introduce a new, label-free (LF) implementation of this approach: FLEXIQuant-LF.

FLEXIQuant-LF enables unbiased identification of differentially modified peptides and quantification of the modification extent in label-free mass-spectrometry data. Our tool only requires raw peptide intensities as input data making it widely applicable. FLEXIQuant-LF determines differentially modified peptides using RANSAC-based robust linear regression in combination with a sophisticated normalization and score calculation procedure (**Fig. 1B**). The RANSAC algorithm iteratively determines outliers in the data and fits the regression line only to observations not classified as outliers (inliers). This ensures that the regression line is fitted to unmodified peptides and facilitates the quantification of the modification extent of modified peptides based on the distance to the regression line.

We benchmarked FLEXIQuant-LF by analyzing the anaphase promoting complex/cyclosome (APC/C) during mitotic arrest. Application of FLEXIQuant-LF to this complex for proof of concept had two advantages: 1) excellent enrichment protocols for isolation of the complex by targeting the cell division cycle protein 27 (CDC27) are well established (17) and 2) the cell cycle-dependent phosphorylation dynamics of APC/C have been qualitatively and quantitatively described (17) (25). We studied the APC/C after treating thymidine-synchronized HeLa cells with nocodazole for 4h, 8h and 10h to induce prometaphase arrest. The APC/C isolated from S-phase cells served as a control (workflow in **Fig. 1A**).

The software is open source and available in two versions: an easy-to-use graphical user interface (GUI) version as well as a command line interface (CLI) version to allow for integration into bioinformatics analysis pipelines.

### LC/MS Analysis of the APC/C

Prior to analysis of the modification dynamics of the APC/C, the complex was purified from each time point separately by co-immunoprecipitation (co-IP) using an anti-CDC27 antibody (workflow in **Fig. 1A**). The protein digests of the co-immunoprecipitates from the different cell cycle stages were analyzed in triplicate using an LC-SWATH method. The SWATH dataset was analyzed using a project specific spectral library comprising 13,724 peptides representing 2,041 proteins (details in *Material and Methods*). Two out of 12 technical replicates were excluded from the analysis due to low number of identified/quantified proteins (one each from the 4h and 10h time points). As the APC/C is a well characterized protein complex, we focused on the bona fide members of this protein complex, namely APC1, APC4, APC5, APC7, APC10, APC15, APC 16, CDC16, CDC20, CDC23, CDC26 and CDC27 precision (raw intensities of all peptides can be found in **Supplementary Table 1**). For further FLEXIQuant-LF analysis, only unmodified peptides, and peptides with methionine oxidation or carbamidomethylation on cysteine were considered. Furthermore, any peptide had to be reliably quantified across all runs. In total, 248 out of 289 APC/C component-derived peptides (86%) met those criteria. Across these 248 quantified peptides, the median Pearson correlation coefficient between replicates was 0.98 (range: 0.96 to 0.99) highlighting excellent quantification.

### Benchmarking FLEXIQuant-LF *with CDC27 and APC5*

For the benchmarking of FLEXIQuant-LF, we focused on CDC27 and APC5. Cell cycle-dependent modification dynamics have been extensively studied for both proteins (17) (25). These pre-existing published data enabled us to validate our peptide classifications.

We quantified 33 peptides corresponding to CDC27 (sequence coverage: 55%), of which nine peptides were classified as likely differentially modified within the time series (**Fig. 2A**, magenta bars) by our FLEXIQuant-LF method. Four peptides were classified as possibly differentially modified (**Fig. 2A**, blue bars). One peptide (_31_LYAEVHSEEALFLLATCamYYR_50_) was classified as outlier based on its raw scores and removed before RM score calculation.

**Figure 2.**
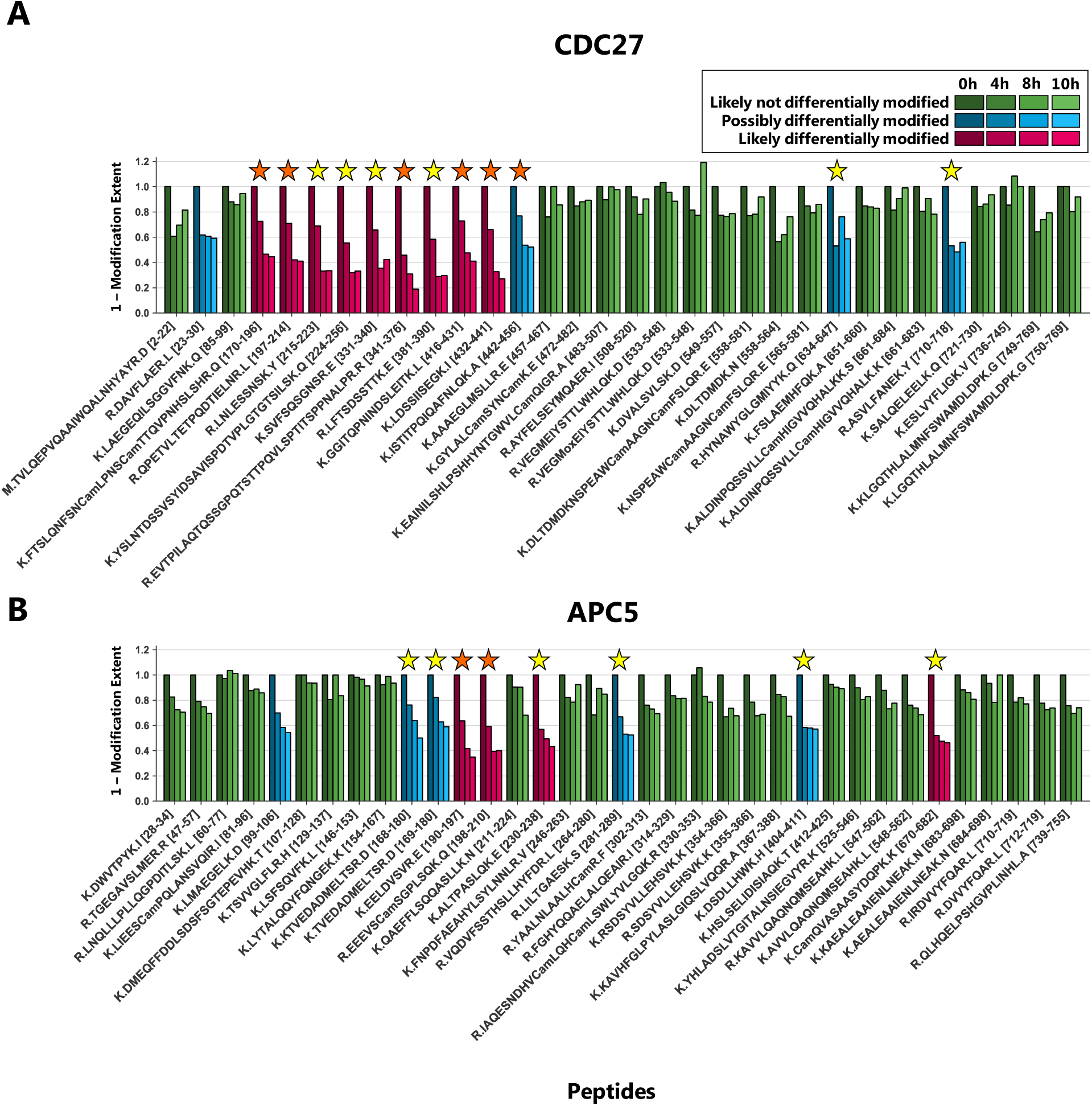
Benchmarking FLEXIQuant-LF with CDC27 and APC5. Peptide classification and extent of modification quantification of CDC27 peptides (A) and APC5 peptides (B). Green bars indicate peptides that were classified as likely <u>not</u> differentially modified (RM score ≥ 0.6), blue bars indicate peptides classified as possibly differentially modified (0.5 ≤ RM score < 0.6) and magenta bars indicate peptides that were classified as likely differentially modified (RM score ≤ 0.5). Shading indicates timepoints from 0h (S phase; darkest shade) to 10h (brightest shade). Positions of peptides within the protein are given from the N- to C-terminus. Orange stars indicate modifications identified in the DDA dataset while yellow stars indicate modifications described in the literature. **A)** FLEXIQuant-LF analysis of 32 CDC27 peptides after filtering classified nine and four peptides as likely and possibly differentially modified over the course of the experiment. We found PTM evidence for all peptides classified as likely differentially modified (for five in our DDA data and for all peptides online (see also **Table 1**)) as well as for three out of four peptides classified as possibly differentially modified (for one in our DDA data and all except of peptide _23_DAVFLAER_30_ described online). **B)** Out of 35 quantified peptides of APC5, four and five peptides were classified as likely and possibly differentially modified. We found evidence for the likely differentially modified peptides in our DDA data or as previously reported. Additionally, we found evidence for four out of five possibly differentially modified peptides online. Only for _99_LMAEGELK_106_, we could not find any evidence for modification.

We first checked for modified peptide species within our DDA dataset. In total, we identified six CDC27-derived phosphorylated peptides in the DDA data (**Fig. 2A**, orange stars). The FLEXIQuant-LF analysis classified the unmodified versions of these six phosphopeptides as follows: five likely differentially modified peptides and one possibly differentially modified peptide. For the other four peptides identified as likely differentially modified as well as for two out of the three remaining peptides classified as possibly differentially modified by FLEXIQuant-LF, phosphorylations and/or ubiquitinylations were described (25) or deposited online at www.phosphosite.org (41) (yellow stars). In summary, FLEXIQuant-LF demonstrated an excellent performance in detecting potentially modified peptides. We found evidence of modification for all peptides identified as likely modified in our study and for three out of four possibly differentially modified peptides.

In addition to the detection of potentially modified peptides, we designed FLEXIQuant-LF to also quantify the degree of differential modification. Relative to time point 0h (S-phase) in our time series, the N-terminal region of CDC27 (amino acids 170-441) was increasingly modified after 4h (27% to 54%, average: 36%) and reached its maximum modification extent after 10h (between 55% and 81%, average: 65%) (**Fig. 2A**). The peptide spanning amino acids 442 to 456 (_442_ISTITPQIQAFNLQK_456_) also showed an increasing degree of modification but to a lower extent (up to 48% modified after 10h).

For APC5, two peptides (_497_FPPNSQHAQLWMLCamDQK_513_ and _35_IAVLVLLNEMSR_46_) out of 37 quantified peptides (sequence coverage: 59%) were classified as outliers based on their raw scores and were excluded from further analysis. Out of the 35 remaining peptides, four were classified as likely differentially modified across the time points (**Fig. 2B**, magenta bars), while five peptides were considered as possibly differentially modified (**Fig. 2B**, blue bars). We identified in our DDA data the phosphorylated cognates for two of the four likely modified peptides (**Fig. 2B**, orange stars). We found evidence of modifications within the literature for the other two peptides identified as likely differentially modified in our FLEXIQuant-LF analysis as well as for four of the five peptides classified as possibly differentially modified (yellow stars) (41). Interestingly, we did not find evidence for modification for one possibly differentially modified peptide (_99_LMAEGELK_106_). This could be interpreted either as a false positive classification or that this peptide has a hitherto undescribed modification on the methionine, glutamic acid or lysine residues.

Overall, these findings are in excellent agreement with the original label-based FLEXIQuant analysis of CDC27 and APC5 (17), which identified the same protein regions as being modified to a comparable extent. It was noted that compared to the original FLEXIQuant study the extent of modification was lower in our novel FLEXIQuant-LF analysis. A highly probable explanation for this phenomenon could be that there was a basal modification level of up to 20% for some peptides in S-phase, as shown with FLEXIQuant (17) which determines the absolute extent of modification. Thus, we emphasize that the FLEXIQuant-LF approach provides the extent of modification relative to the given reference time point.

### The modification dynamics of other APC/C complex components

FLEXIQuant-LF has an advantage over the conventional FLEXIQuant implementation in that the analysis can easily be performed on all proteins within a sample (provided a minimum of five peptides was quantified to allow for linear regression). Here, we extended our analysis to all other APC/C proteins of which sufficient peptides were reliably quantified in our data, namely APC1 (**Fig. 3**), APC4, APC7, APC10, CDC16, CDC20 and CDC23 (**Fig. 4**, individual figures for each protein including the peptide sequences can be found in **Supplementary Fig. 1**). By applying FLEXIQuant-LF, we classified between zero (APC4, APC10) and seven peptides (APC1) per protein as likely differentially modified and between zero (APC4, APC10, CDC20) and four peptides (APC1) per protein as possibly differentially modified (summary in **Table 1** and **Table 2**).

**Table 1.** Overview of FLEXIQuant-LF analysis for the APC/C complex components during nocodazole arrest; quant. peptides: quantified peptides; seq. cov.: sequence coverage; likely diff. modified: number of peptides classified as likely differentially modified; possibly diff. modified : number of peptides classified as possibly differentially modified; excluded: number of peptides classified as outliers and thus excluded from RM score calculation; DDA: number of peptides classified as differentially modified for which we found evidence in our DDA data; lit.: number of peptides classified as differentially modified for which we found evidence for modification in the literature).

| protein | quant.<br>peptides | seq.<br>cov. | FLEXIQuant-LF analysis |  |  | evidence |  |  |
| --- | --- | --- | --- | --- | --- | --- | --- | --- |
|  |  |  | likely<br>diff.<br>modified | possibly<br>diff.<br>modified | ex-<br>cluded | DDA | lit. | no<br>evidence |
| APC1 | 54 | 39% | 7 | 4 | 4 | 8 | 9 | 2 |
| APC4 | 28 | 43% | 0 | 0 | 2 | - | - | 0 |
| APC5 | 37 | 59% | 4 | 5 | 2 | 2 | 8 | 1 |
| APC7 | 27 | 52% | 4 | 2 | 1 | 1 | 4 | 2 |
| APC10 | 9 | 69% | 0 | 0 | 3 | - | - | 0 |
| APC15 | 1 | 30% | 0 | 0 | 0 | - | - | 0 |
| APC16 | 3 | 30% | 0 | 0 | 0 | - | - | 0 |
| CDC16 | 23 | 41% | 3 | 2 | 3 | 1 | 3 | 2 |
| CDC20 | 7 | 22% | 1 | 0 | 1 | 0 | 1 | 0 |
| CDC23 | 33 | 52% | 3 | 3 | 4 | 3 | 5 | 1 |
| CDC27 | 33 | 55% | 9 | 4 | 1 | 6 | 12 | 1 |
| ***in total*** | **248** | **AVG**<br>**47%** | **30**<br>**12%** | **20**<br>**8%** | **20**<br>**8%** | **21**<br>**42%** | **41**<br>**82%** | **9**<br>**18%** |

**Table 2.**
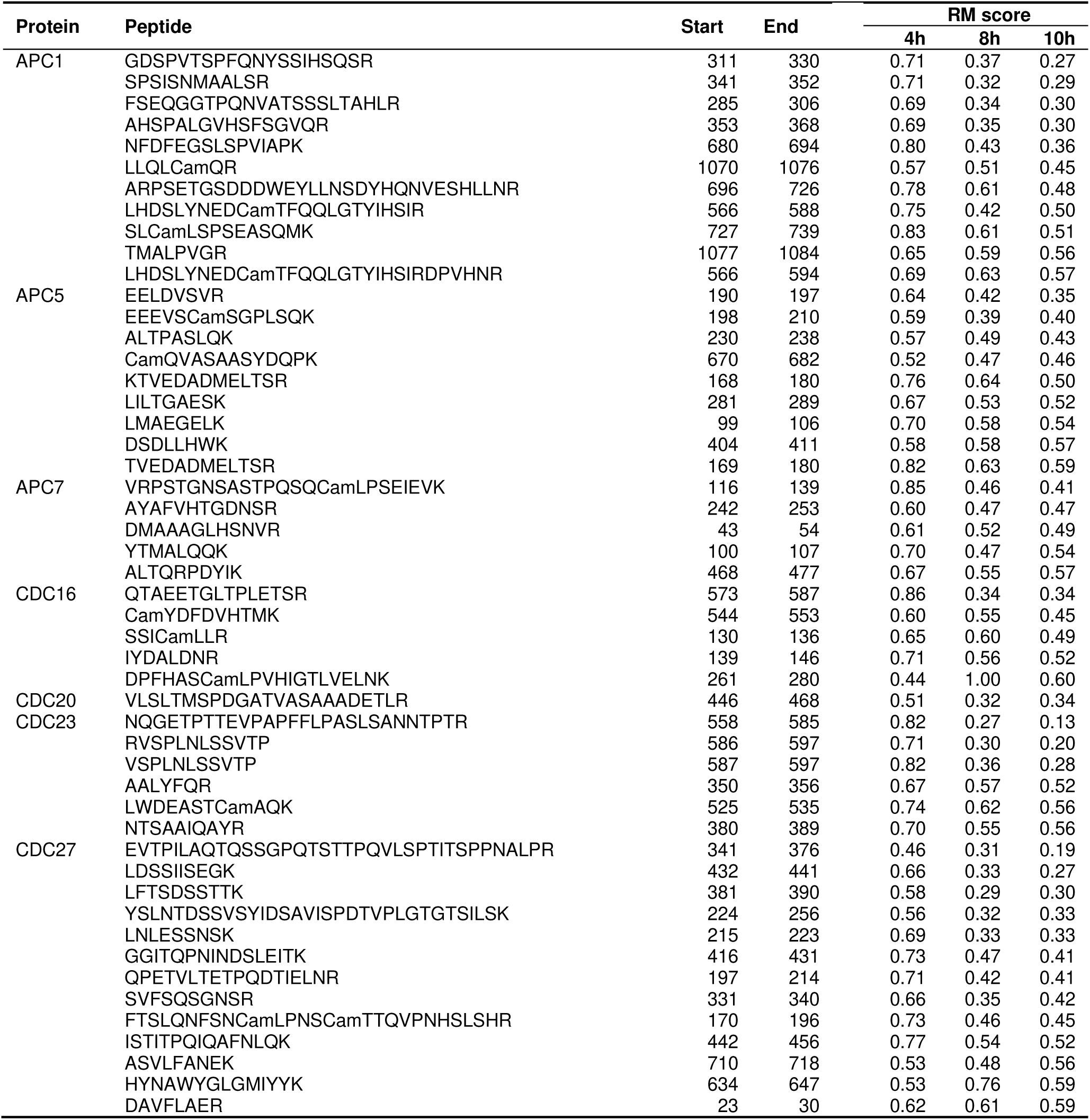
Differentially modified peptides. Overview of peptides classified as likely or possibly differentially modified and resulting RM scores of the FLEXIQuant-LF analysis of APC/C.

**Figure 3.**
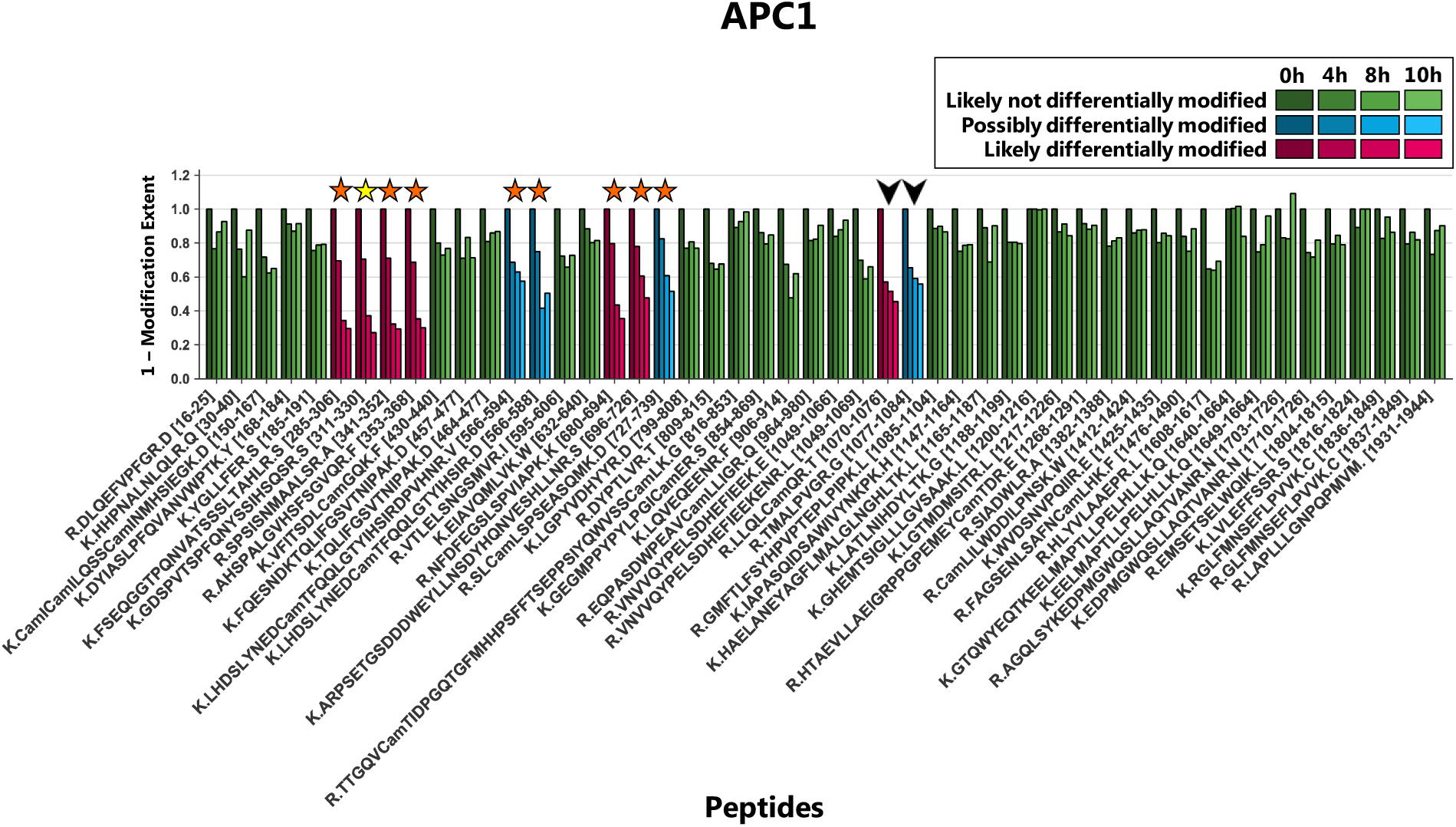
Application of FLEXIQuant-LF to APC1. Peptide classification and extent of modification of APC1 (see **Figure 2** for details about colors and shading). Seven out of 50 quantified peptides were classified as likely differentially modified and four peptides as possibly differentially modified. We found evidence of modification for six out of seven peptides classified as likely differentially modified (five in our DDA data and six described in the literature) as well as for three out of four peptides classified as possibly differentially modified in our DDA data (see also **Table 1**). For one peptide classified as likely differentially modified (_1070_LLQLCamQR_1076_) and for one peptide classified as possibly differentially modified (_1077_TMALPVGR_1084_), we could not find evidence of modification. Interestingly, the two peptides (indicated by arrow heads) are consecutive peptides and the second peptide starts with threonine (AA 1077). An as of now undiscovered threonine modification such as a phosphorylation would likely lead to a highly increased missed cleavage rate and could explain the reduction of the signal intensities of both peptides.

**Figure 4.**
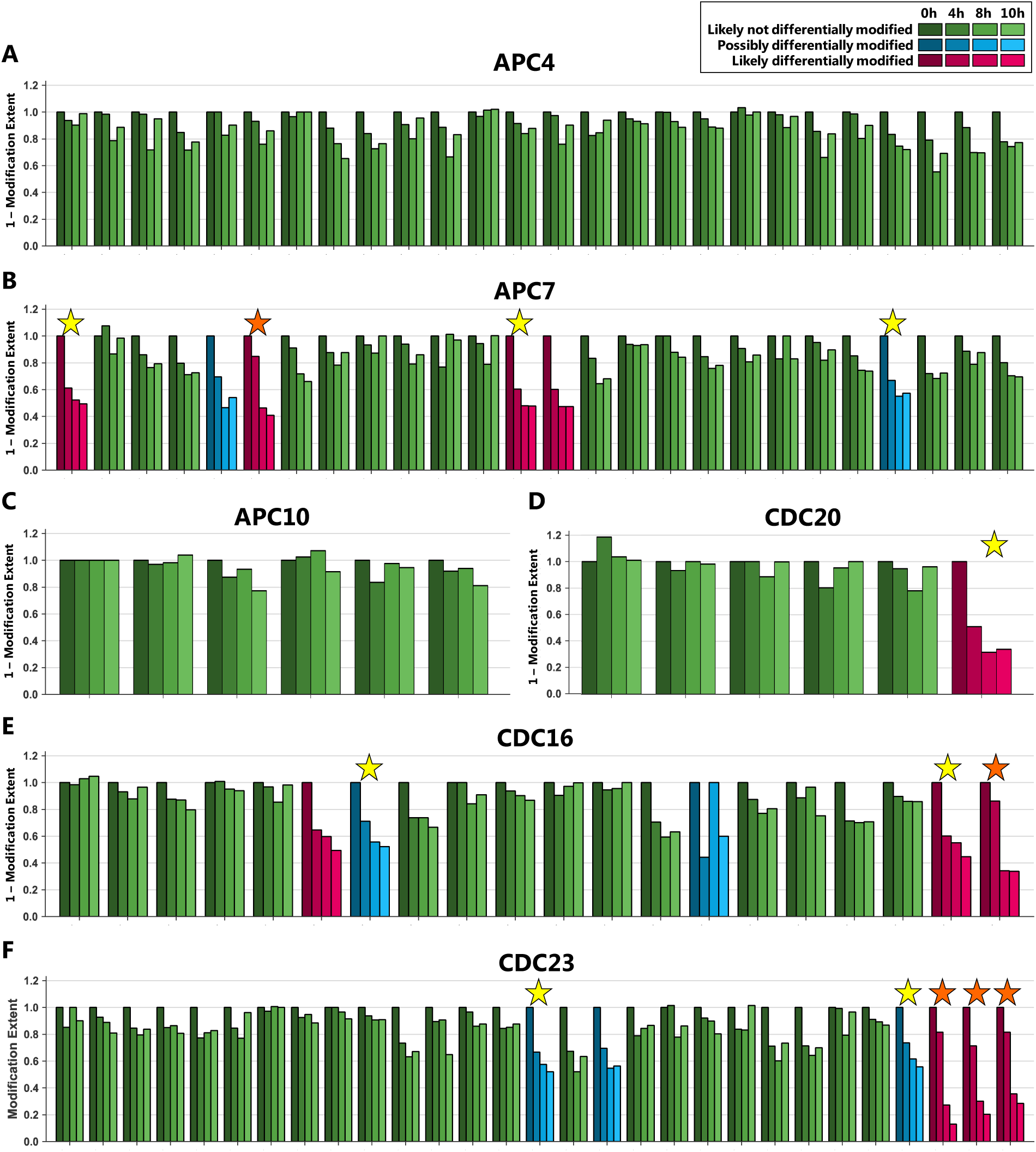
Application of FLEXIQuant-LF to the remaining APC/C core components. Peptide classification and extent of modification of **A:** APC4, **B:** APC7, **C:** APC10, **D**: CDC20, **E:** CDC16 and **F:** CDC23. See **Figure 2** for color and shading scheme.

From APC1, we quantified 54 peptides (sequence coverage: 39%) in total, of which four peptides were classified as outlier based on their raw scores and were excluded from further analysis (_930_LVVWMTNVGFTLR_942, 943_DLETLPFGIALPIR_956, 1546_TGGEMNYGFHLAHHMALGLLFLGGGR_1571_ and _1618_LLVPVDVDTNTPCamYALLEVTYK_1639_). Out of the remaining 50 peptides, seven peptides were classified as likely differentially modified and four peptides as possibly differentially modified (**Fig. 3**, magenta and blue bars respectively). We identified phosphorylated peptides in the DDA data for five of the seven likely and for three out of four possibly differentially modified peptides (**Fig. 3**, orange stars).

We also classified the peptide _311_GDSPVTSPFQNYSSIHQSR_330_ as likely differentially modified by our FLEXIQuant-LF analysis, although no modified cognate peptides were identified in the DDA data. However, multiple modifications, i.e. phosphorylations on S313, T316, S317, Y322 and S324, have been reported in previous studies (41) (indicated by a yellow star in **Fig. 3**). For one likely (_1070_LLQLCamQR_1076_) and one possibly differentially modified peptide (_1077_TMALPVGR_1084_), we could not find published evidence of PTMs within the sequence of those two peptides. Interestingly, these are consecutive peptides with the latter one starting with a threonine residue (AA 1077). A hitherto undiscovered phosphorylation at this threonine residue could lead to a highly increased missed cleavage which would be consistent with the observed intensity reduction of both peptides.

A more detailed analysis of the modification dynamics observed in APC1 identified two regions within the protein that appear to have different modification kinetics (**Fig. 3**). Peptides in the N-terminal part ranging from residues 285-368 (4 peptides), showed a degree of modification of 29-31% after 4 hours, 63-68% after 8 hours and between 70% and 71% after 10 hours. In contrast, the peptides in a more C-terminal domain spanning residues 680-739 (3 peptides) showed slower modification kinetics and a lower degree of modification at 10 hours. After 4 hours, 17-22% change in modification status was observed. The extent of modification changed between 39% and 57% after 8 hours, and 49-64% after 10 hours. These data demonstrate that tools to study the modification dynamics in a quantitative manner will greatly improve the current understanding of biological processes.

### Analyzing multiple proteins as a “superprotein”

For the two smallest APC/C subunits, namely APC15 and APC16, the number of reliably quantified peptides were 1 and 3, respectively, i.e. below the described threshold of 5 quantifiable peptides, making the above described FLEXIQuant-LF analysis nonapplicable. However, if proteins are known to have invariable expression/abundance profiles over the considered time period or between different conditions of interest, as e.g. observed for the core components of the APC/C, we hypothesized that the peptides of multiple proteins can be analyzed together by treating this set of proteins with invariable abundance as one “superprotein”. This would expand the FLEXIQuant-LF approach to smaller proteins and allow for the identification and quantification of differential modification of proteins with less than five quantified peptides that could not be analyzed by the normal FLEXIQuant-LF implementation otherwise.

To test this “superprotein” approach, we applied FLEXIQuant-LF to analyze all reliably quantified APC/C proteins in our data in a combined fashion as a “superprotein”, namely APC1, APC4, APC5, APC7, APC10, APC15, APC16, CDC16, CDC23 and CDC27. CDC20 was excluded from this “superprotein” approach as it is known that this APC/C regulator displays changing affinity to the APC/C during the M-phase of the cell cycle, i.e. its abundance within the APC/C changes during the investigated time period (42). We validated the “superprotein” strategy by comparing its results to the results of the FLEXIQuant-LF analysis of the individual proteins for all proteins that could be analyzed with both strategies. The two approaches resulted in identical classification of peptides as well as near identical quantification of the modification extent (mean RM score difference = 2.45E-10, r^2^ = 1), demonstrating the applicability of the “superprotein” strategy (**Supplementary Table 2**).

The FLEXIQuant-LF analysis of APC15 and APC16 using the “superprotein” approach revealed that all four peptides associated with these two small proteins not amenable to the conventional FLEXIQuant-LF approach are likely not differentially modified (**Supplementary Table 2** and **Supplementary Fig. 1 F and G**). These results are in accordance with the literature where, to the best of our knowledge, no cell cycle-dependent differential modifications in APC15 and APC16 have been described for the detected protein domains (41).

## Discussion

Our FLEXIQuant-LF is based on robust linear regression, as implemented in the RANSAC algorithm. In general, linear regression requires a certain number of data point to yield meaningful results and becomes more accurate with increasing number of data points. Thus, FLEXIQuant-LF becomes more powerful with increasing numbers of unique peptides that can be identified for the proteins of interest. We set the minimum number of unique peptides required for a FLEXIQuant-LF analysis to five, although a larger number of peptides will provide more robust results. Addressing this limitation, we introduce the “superprotein” approach.

In the presented proof-of-concept study, we applied FLEXIQuant-LF to a time series experiment. However, FLEXIQuant-LF can also be applied to study protein modification differences between different experimental conditions or cohorts in general, e.g. comparing diseased vs. healthy individuals or testing the effect of different drugs. As FLEXIQuant-LF only determines the extent of differential modification of a peptide relative to the reference, quantification of the absolute abundance of the modification still requires an internal reference. However, using a reference time point or condition has the advantage of highlighting modifications that exhibit dynamic regulation in the context of interest. Furthermore, FLEXIQuant-LF can be used to repeatedly analyze the same data set with varying reference samples, providing complementary perspectives on the modification dynamics. This consideration is particularly important for the current implementation of FLEXIQuant-LF which only quantifies modifications extents that are increasing relative to the reference sample. This is due to our normalization method used for RM score calculation, which excludes peptides with decreased modification. FLEXIQuant-LF reports these peptides in the “_removed_peptides.csv” output file and raw score information can be retrieved from the “_raw_score.csv” output file. We reasoned that the reference sample is normally the least modified protein. In case this assumption does not hold, a second FLEXIQuant-LF can be carried out after selecting another samples as the reference.

While FLEXIQuant-LF, similar to the initial FLEXIQuant approach, identifies peptides that are differentially modified, the specific type and site of modification is not the objective of this analysis. The detailed localization and characterization of the modifications require either more in-depth data analysis or follow-up experiments. Nevertheless, this analysis method can be easily applied in any affinity purification or global proteomics experiment and the results can further be used to identify proteins to be synthesized as reference standards, if more in-depth experiments are needed. In the original FLEXIQuant approach, changes in the modification extent had to be above 20% in order to be reliably detected. For our proof of concept study, we chose 40% which is consistent with stoichiometries commonly observed in cell cycle-focused or signaling pathway activation-related large scale phosphoproteomics studies, which reported serine and threonine phosphorylation extents exceeding 75% and tyrosine phosphorylation extents above 50% (43). However, our FLEXIQuant-LF approach enables the selection of a minimum change in modification extent deemed to be interesting and/or compatible with the precision of the dataset; clearly the usual trade-off considerations between specificity and sensitivity apply.

Even though APC/C is a moderately abundant complex within the cell, we opted for affinity enrichment to ensure a high number of unique peptides per protein of the complex components are measured. For highly abundant protein classes such as the ribosome and proteasome (44), a direct FLEXIQuant-LF analysis on the whole cell lysate is most likely a viable option. However, we expect that future technological developments in LC/MS instrumentation and data analysis approaches will make less abundant proteins and protein complexes such as signaling proteins/complexes amenable to FLEXIQuant-LF analysis without further enrichment of the proteins/protein complexes of interest.

A premise of the FLEXIQuant-LF method is, that the modification state of the majority of peptides in a protein does not change between experimental conditions or over the time period considered in a study. This is likely to hold true for most proteins. However, in some cases where large parts of a protein are differentially processed and/or modified, as in Tau for example (22), FLEXIQuant-LF might not be applicable.

The combination of affinity purification of the protein complex and DIA analysis facilitated the precise and reproducible peptide quantification. Currently, DIA is the most suitable acquisition method due to its precise peptide quantification capabilities with fewer missing values. However, the FLEXIQuant-LF approach is also applicable to other label-free quantification methods.

## Conclusions

Here, we presented FLEXIQuant-LF, a computational label-free implementation of the original FLEXIQuant method using a DIA dataset of the APC/C during mitotic arrest as a proof of concept. FLEXIQuant-LF is a widely applicable analysis workflow building on the previously published FLEXIQuant idea of monitoring the unmodified peptides and changes to their signal intensities to detect modification events and to quantify the degree of modifications. The use of a reference timepoint or reference sample instead of labeled reference proteins allows FLEXIQuant-LF to be universally applicable post hoc to any protein robustly identified across the sample series without the need for the addition of heavy isotope-labeled protein standard(s).

The application of FLEXIQuant-LF to a DIA dataset of the APC/C isolated from Hela cells arrested at various time points during a mitotic arrest, allowed us to recapitulate the modification dynamics previously observed with our original heavy isotope-labeled internal standard-based FLEXIQuant method. Using FLEXIQuant-LF, we easily extended our analysis to other APC/C subunits present in the sample set and delivered a comprehensive report on the modification dynamics of the core components of APC/C following nocodazole-induced mitotic arrest. Furthermore, we introduced a “superprotein” approach which allowed us to include proteins, which provided an insufficient numbers of unique peptides to be analyzed individually; this is a limitation that applies in particular to small proteins, but can also be encountered in very acidic or hydrophobic proteins with few proteolytic cleavage sites. This “superprotein” strategy enabled us to extract more information from data and gain additional insights about peptides/proteins that would have been disregarded otherwise.

FLEXIQuant-LF is unbiased towards the type of modification and relies on identification of the unmodified counterpart of a modified peptide. This valuable information can subsequently be used to study rare PTMs and to potentially discover novel PTMs using more targeted approaches in future experiments. Therefore, FLEXIQuant-LF can pave the way to a better general understanding of post-translational modification dynamics of proteins. We expect that FLEXIQuant-LF will enable researchers to revisit their data and identify novel modification-driven features of health and disease that could be used as targets for treatment.

## Supporting information

Supplementary Reproducibility

## Abbreviations

AP: Affinity purification
APC/C: Anaphase promoting complex/cyclosome
CLI: Command line interface
Co-IP: Co-immunoprecipitation
CV: Coefficient of variance
DDA: Data-dependent acquisition
DIA: Data-independent acquisition
FLEXIQuant: Full-length-expressed stable isotope-labeled proteins for quantification
GUI: Graphical user interface
LF: label-free
MAD: Median absolute deviation
PTM: Post translational modification
RANSAC: Random sample consensus
RM score: Relative modification score
SRM: Selected reaction monitoring
SWATH: Sequential window acquisition of all theoretical fragment-ion spectra

## Data and Software Availability

Data are available via ProteomeXchange with identifier PXD018411 (Reviewer access with username: and password: BWxUSwIR). FLEXIQuant-LF is available with graphical user interface (GUI) as standalone executable EXE file. Additionally, a command line interface (CLI) version is available as PY file. Both versions as well as the Python source code can be downloaded via GitHub: https://github.com/SteenOmicsLab/FLEXIQuantLF.

## Acknowledgements

We acknowledge the following funding from the US National Institutes of Health: S10OD0107060 to H.S. for the TripleTOF 5600 mass spectrometer, R01CA196703, R01AI099204 and U01AI124284 covering fractions of the efforts from H.S., R01NS066973 covering R.C.’s effort, RC4GM096319 covering J.M.’s effort and R01GM112007 to J.A.S. covering part of her and C.N.S.’ effort. B.Y.R. gratefully acknowledges financial support from Deutsche Forschungsgemeinschaft (RE3474/2-2) covering part of his and K.K.’s effort. We thank Benoit Fatou and Mukesh Kumar for providing test data during development and testing of FLEXIQuant-LF. Furthermore, we thank Kyle Higgins and Patrick van Zalm for testing the FLEXIQuant-LF software.

## Author contributions

Conceptualization, H.S., J.A.S., B.Y.R. C.N.S.; Methodology K.K., C.N.S., B.Y.R., H.S.; Software, K.K.; Validation, K.K., C.N.S., H.S., B.Y.R.; Formal Analysis, K.K.; Investigation, J.M., R.C., K.K.; Resources: H.S., J.A.S.; Data Curation, K.K., C.N.S.; Writing – Original Draft, K.K., J.M.; Writing – Review & Editing, C.N.S., H.S., B.Y.R.; Visualization, K.K., J.M.; Supervision, H.S., B.Y.R, C.N.S.; Project Administration, H.S.; Funding Acquisition, H.S., B.Y.R., J.A.S.;

## Supporting Information

**Supplementary Figure 1.** FLEXIQuant-LF analysis results of all APC/C components. **A:** APC1, **B:** APC4, **C:** APC5, **D:** APC7, **E:** APC10, **F:** APC15, **G:** APC16, **H:** CDC16, **I:** CDC20, **J:** CDC23, **K:** CDC27.

**Supplementary Table 1.** Raw peptide intensities of all identified APC/C components

**Supplementary Table 2.** Comparison RM scores proteins analyzed individually and using the “superprotein” approach.

**Supplementary Methods**

**Supplementary Reproducibility**

