## Supplementary Methods for "FLEXIQuant-LF: Robust Regression to Quantify Protein Modification Extent in Label-Free Proteomics Data"

### Supplementary Information

#### Detailed methods

##### Derivation of RM Score

FLEXIQuant-LF is based on the assumption that the intensities of the peptides of a protein of interest are linearly proportional to the intensities of their counterparts in a reference sample and the distance of each peptide to the regression line is measured. Then these distances would correlate with the degree of deviation from the reference sample but would not be an absolute measure of it. Three factors influence the distance to the regression line in this scenario:

- Extent of modification of a peptide
- Concentration of a protein in a sample
- Intensity of a peptide

To demonstrate the influences of each factor we created small artificial data sets where only one factor is altered at a time.

**Influence of the Extent of Modification of a Peptide**

The basic principle of the developed quantification method, namely determining the degree of modification based on the distance to the regression line, can be easily illustrated with the artificial data set shown in Table 1. All peptides in sample 1 are unmodified, while in sample 2 - sample 4 the intensity of peptide 5 is decreased by 25%, 50% and 75%, respectively, simulating differing degrees of modification. The linear regression plot (Figure 1) demonstrates that the distance to the regression line increases with an increasing extent of modification.

*Table 1: Artificial data demonstrating the influence of the extend of modification of a given peptide. All peptides in sample 1 are unmodified, while in sample 2 - sample 4 the intensity of peptide 5 is decreased by 25%, 50% and 75%, respectively.*

| Peptide | Reference | Sample 1 | Sample 2 | Sample 3 | Sample 4 |
|---|---|---|---|---|---|
| Peptide 1 | 100 | 100 | 100 | 100 | 100 |
| Peptide 2 | 200 | 200 | 200 | 200 | 200 |
| Peptide 3 | 300 | 300 | 300 | 300 | 300 |
| Peptide 5 | 400 | 400 | 300 | 200 | 100 |
| Peptide 6 | 500 | 500 | 500 | 500 | 500 |

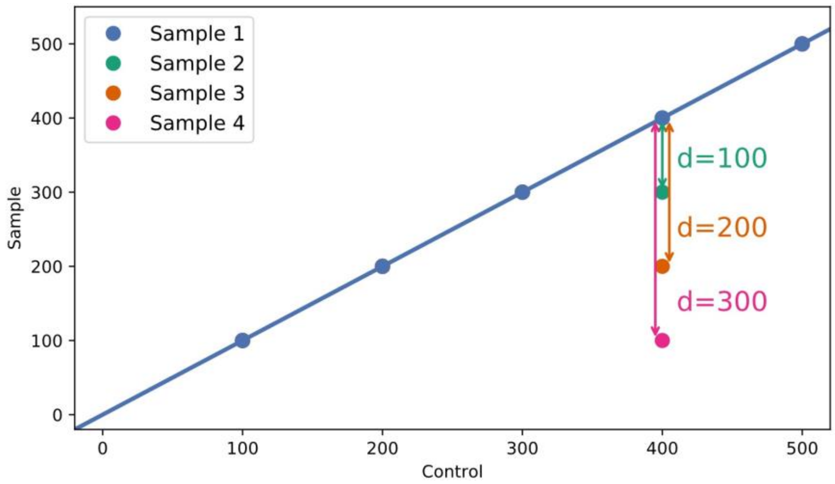


*Figure 1: Exemplified influence of extent of modification on the distance to regression line. Sample 1 is unmodified, while in samples 2-4 the intensity of peptide 5 is reduced by 25%, 50% and 75%,respectively, compared to Sample 1, demonstrating the influence of the extent of modification of a peptide on the distance.*

##### Influence of the Concentration of a Protein in a Sample

The slope of the regression line in the regression plots is dependent on the concentration of the observed protein. If the reference samples remain unchanged the slope will increase with increasing concentration of a given protein. In the artificial data set in Table 2 the intensities of sample 2 are three times as high as in sample 1. In both samples the intensity of peptide 3 is decreased by 50% compared to the predicted intensity (point on the regression line at x=300). The slope of the regression line of sample 1 is 0.5 whereas the slope of sample 2 is 1.5. Figure 2 demonstrates that the distance of peptide 3 to the regression line is different for sample 1 (d=75) than for sample 2 (d=225) even though both intensities were decreased by 50%. To enable the comparison between samples with different concentrations of the protein of interest, a normalization of the distance that addresses this factor of influence is necessary. A simple way to achieve this is to divide the distance by the slope:

Sample 1: $\frac{75}{0.5} = 150$ Sample 2: $\frac{225}{1.5} = 150$

$$normalized\ distance = \frac{distance\ to\ regression\ line}{slope\ of\ regression\ line} \quad (1)$$

*Table 2: Artificial data demonstrating the influence of the concentration of a protein in a sample. The intensity of peptide 3 is decreased by 50% in sample 1 and sample 2, while the overall abundance of peptides in sample 2 over sample 1 is 3-fold higher.*

| Peptide | Reference | Sample 1 | Sample 2 |
|---|---|---|---|
| Peptide 1 | 100 | 50 | 150 |
| Peptide 2 | 200 | 100 | 300 |
| Peptide 3 | 300 | 75 | 225 |
| Peptide 5 | 400 | 200 | 600 |
| Peptide 6 | 500 | 250 | 750 |

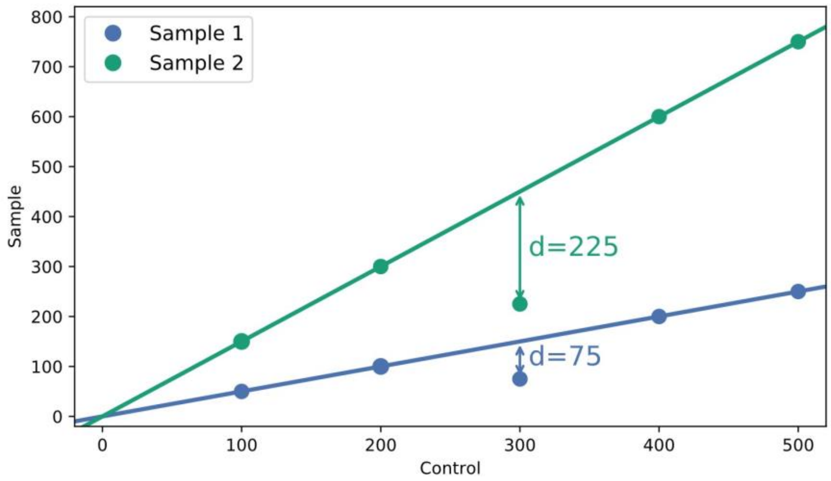


*Figure 2: Influence of protein concentration on the distance to regression line. Peptide 3 is 50% reduced in comparison to the predicted intensity of 150 for sample 1 and 450 for sample 2 with overall abundance difference of 0.5-fold and 1.5-fold of the samples to the reference sample, demonstrating the influence of the concentration on the distance.*

#### Influence of the Intensity of a given Peptide

The intensity of a peptide is another factor that influences its distance to the regression line. Table 3 shows an artificial data set with five samples including one reference sample, one unmodified sample (sample 1) and three samples where the intensity of one peptide per sample is decreased by 50% (sample 2-4). The regression plot (Figure 3) reveals a large difference between the three distances of the modified peptides to the regression line despite each being decreases by the same degree. To account for this influence and thus permit comparison of peptides with different median intensities of the reference samples is to divide the distance by the median intensity of the reference samples.

Sample 2: $\frac{50}{150} = 0.33$ Sample 3: $\frac{150}{450} = 0.33$ Sample 4: $\frac{250}{750} = 0.33$

$$normalized\ distance = \frac{distance\ to\ regression\ line}{instensity\ of\ reference\ samples} \qquad (2)$$

*Table 3: Artificial data set demonstrating the influence of the intensity of a given peptide. Sample 1 is unmodified, in each of the samples 2-4 the intensity of one peptide is decreased by 50% (peptide 1, peptide 3 and peptide 5).*

| Peptide | Reference | Sample 1 | Sample 2 | Sample 3 | Sample 4 |
|---|---|---|---|---|---|
| Peptide 1 | 150 | 100 | 50 | 100 | 100 |
| Peptide 2 | 300 | 200 | 200 | 200 | 200 |
| Peptide 3 | 450 | 300 | 300 | 150 | 300 |
| Peptide 4 | 600 | 400 | 400 | 400 | 400 |
| Peptide 5 | 750 | 500 | 500 | 500 | 250 |

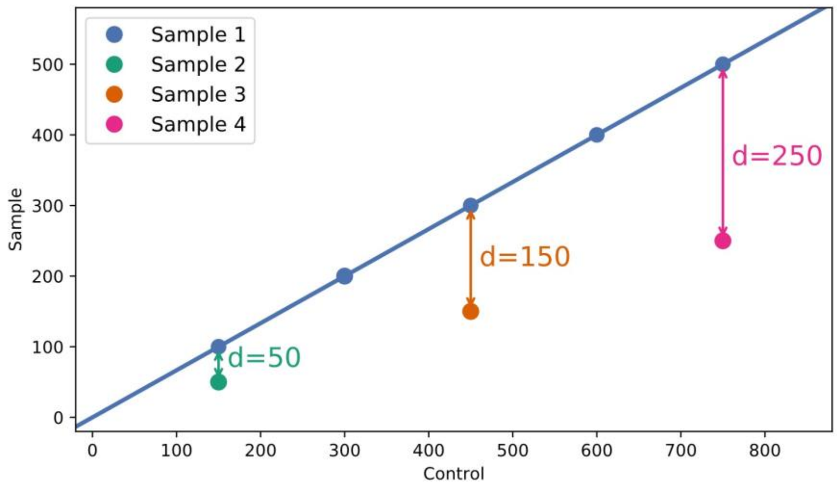


*Figure 3: Influence of peptide intensity on the distance to regression line. Sample 1 is unmodified, in each of the Samples 2-4 the intensity one peptide is 50% reduced compared to Sample 1, demonstrating the influence of the peptide intensity on the distance.*

**RM Score**

Combining the two methods for normalizing the influences of the concentration of a protein in a sample (Equation 1) and the intensity of a given peptide (Equation 2) leads to Equation 3:

$$normalized\ distance = \frac{distance\ to\ regression\ line}{slope \times instensity\ of\ peptide\ in\ reference\ samples} \qquad (3)$$

This is equivalent to dividing the distance to the regression line by the predicted intensity (point on the regression line at the intensity of the peptide in the reference sample):

$$normalized\ distance = \frac{distance\ to\ regression\ line}{predicted\ intensity} \tag{4}$$

The result of this normalization (Equation 4) is the degree of reduction of the peptide intensity compared to the reference, which equals the extent of modification of a given peptide.

In the classical FLEXIQuant method the resulting light/heavy ratio represents the fraction of the unmodified version of a given peptide, meaning a L/H ratio of 1 corresponds to a completely unmodified and 0 to a completely modified peptide. To build this approach analogously the normalized distance which was calculated using Equation 4 is subtracted from 1, yielding the FLEXIQuant-LF raw score:

$$raw\ score = 1 - normalized\ distance = 1 - \frac{distance\ to\ regression\ line}{predicted\ intensity} \tag{5}$$

For each time point, peptides with a raw score larger or equal than three times the median absolute deviation above the median of all raw scores are classified as outliers and excluded from the following score calculation:

$$outlier\!:\ raw\ score\ \geq median(raw\ scores) + 3 \times MAD(raw\ scores) \tag{6}$$

To improve quantification accuracy, raw scores are then scaled for each time point using a “top3” normalization approach. For this, each raw score is divided by the median of the three highest (inlier) raw scores for each time point yielding the final score, termed relative modification (RM) score:

$$RM\ score = \frac{raw\ score}{median(three\ highest\ raw\ scores)} \tag{7}$$
