## Supplementary Reproducibility for "FLEXIQuant-LF: Robust Regression to Quantify Protein Modification Extent in Label-Free Proteomics Data"

### Supplementary Information

#### Reproducibility

The reproducibility of the developed approach was tested using a large, inhouse MRM data set consisting of 81 samples of which 26 were reference group samples. We ran FLEXIQuant-LF 1000 times on the same protein (Tau) and analyzed the resulting slopes to determine the number and frequency of different outcomes. For 6 out of 81 samples the FLEXIQuant-LF yielded ambiguous results. In all six cases two distinct outcomes were observed.

To improve the reproducibility, we initiated the RANSAC algorithm multiple times within a single run (1, 10, 20, 30, 40, 50, 60, 70, 80, 90 and 100) and selected the best model (based on $r^2$ score). This was again tested by running each set of initiations 1000 times and analyzing the slopes of the resulting best models. Initiating RANSAC 10 times resulted in 6 ambiguous samples, 20 initiations in 5, 30 in 2 and 40 in 3. Between 50 and 100 initiations, the number ambiguous samples was oscillating between 1 and 2 (Figure S1 a)). Furthermore, as illustrated by means of sample 30, the frequency of different outcomes within a sample decreased with increasing number of RANSAC initiations (Figure S1 b)). With one initiation the suboptimal outcome prevailed (88%), whereas with 10 initiations the optimal outcome was predominant (67.2%). Using 30 initiations the fraction of suboptimal outcomes was 2.2% and further decreased to 0.5% with 50 initiations.

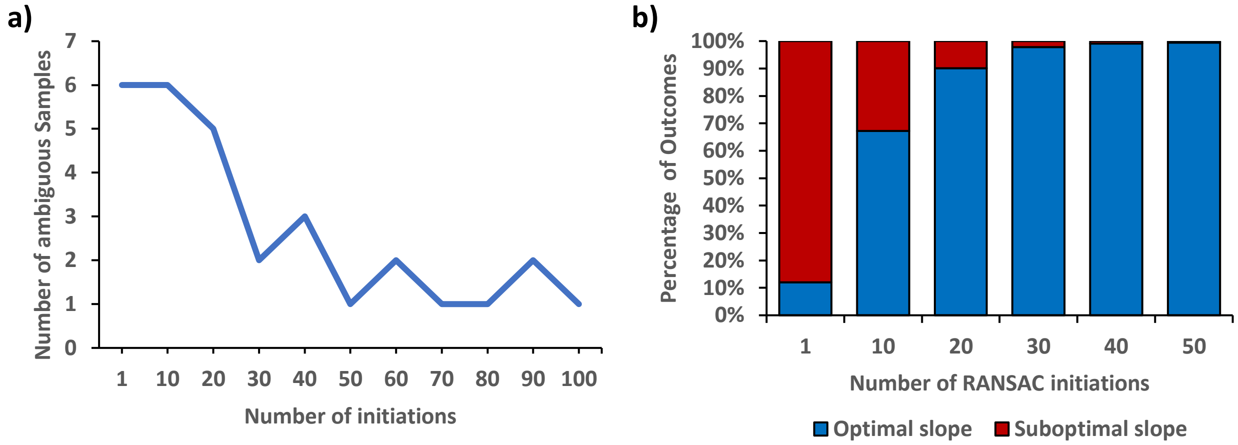


*Figure S1: Reproducibility of PMF. a) Number of ambiguous samples with different number of RANSAC initiations. The number of samples with ambiguous results over 1000 runs decreases with increasing number of RANSAC initiations but starts to oscillate between 1 and 2 ambiguous samples from 50 initiations upwards. b) Fraction of optimal and suboptimal outcomes of PMF with sample 30. The frequency of a suboptimal outcome decreases with increasing number of RANSAC initiations.*

In absolute numbers this means, using 30 RANSAC initiations and selecting the best model resulted in 44 suboptimal outcomes in 81,000 runs of FLEXIQuant-LF (0.05%). These results confirm the benefits of multiple RANSAC initiations and demonstrate a very high reproducibility of the developed approach. Based in these finding the default number of RANSAC initiations in PMF was set to 30 but can be changed by the user.
