## Supplementary Figure 1 for "FLEXIQuant-LF: Robust Regression to Quantify Protein Modification Extent in Label-Free Proteomics Data"

**FLEXIQuant-LF analysis of all APC/C components analyzed as superprotein.**
Peptide classification and extent of modification of . **A:** APC1, **B:** APC4, **C:** APC5, **D:** APC7, **E:** APC10, **F:** APC15, **G:** APC16, **H:** CDC16, **I:** CDC20, **J:** CDC23, **K:** CDC27 analyzed with FLEXIQuant-LF. APC15 and APC16 were analyzed using the “superprotein” approach, the remaining proteins were analyzed individually. Green bars indicate peptides that were classified as likely not differentially modified at 10h (RM score ≥ 0.6), blue bars indicate peptides classified as possibly differentially modified (0.5 ≤ RM score < 0.6) and magenta bars indicate peptides that classified as likely differentially modified (RM score ≤ 0.5). For each class, the four time points are shown in different color shades, from time point 0h (S phase; darkest shade) to time point 10h (brightest shade). Positions of peptides within the protein are given from the N- to C-terminus. Peptides labeled with an orange star indicate that a modification was identified on this peptide in the DDA dataset and peptides labeled with a yellow star indicate a modification that was described in the literature.

## A

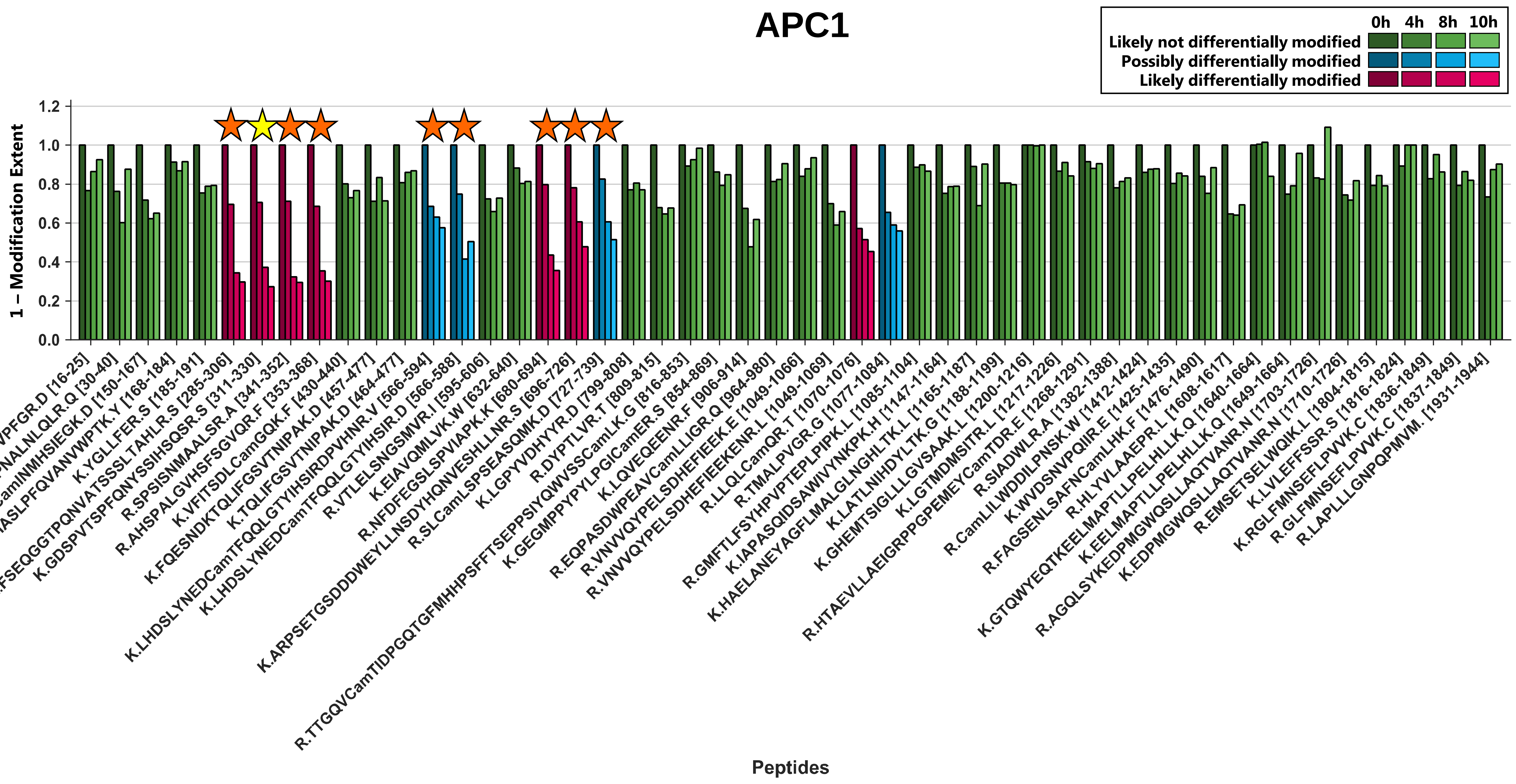

**B**

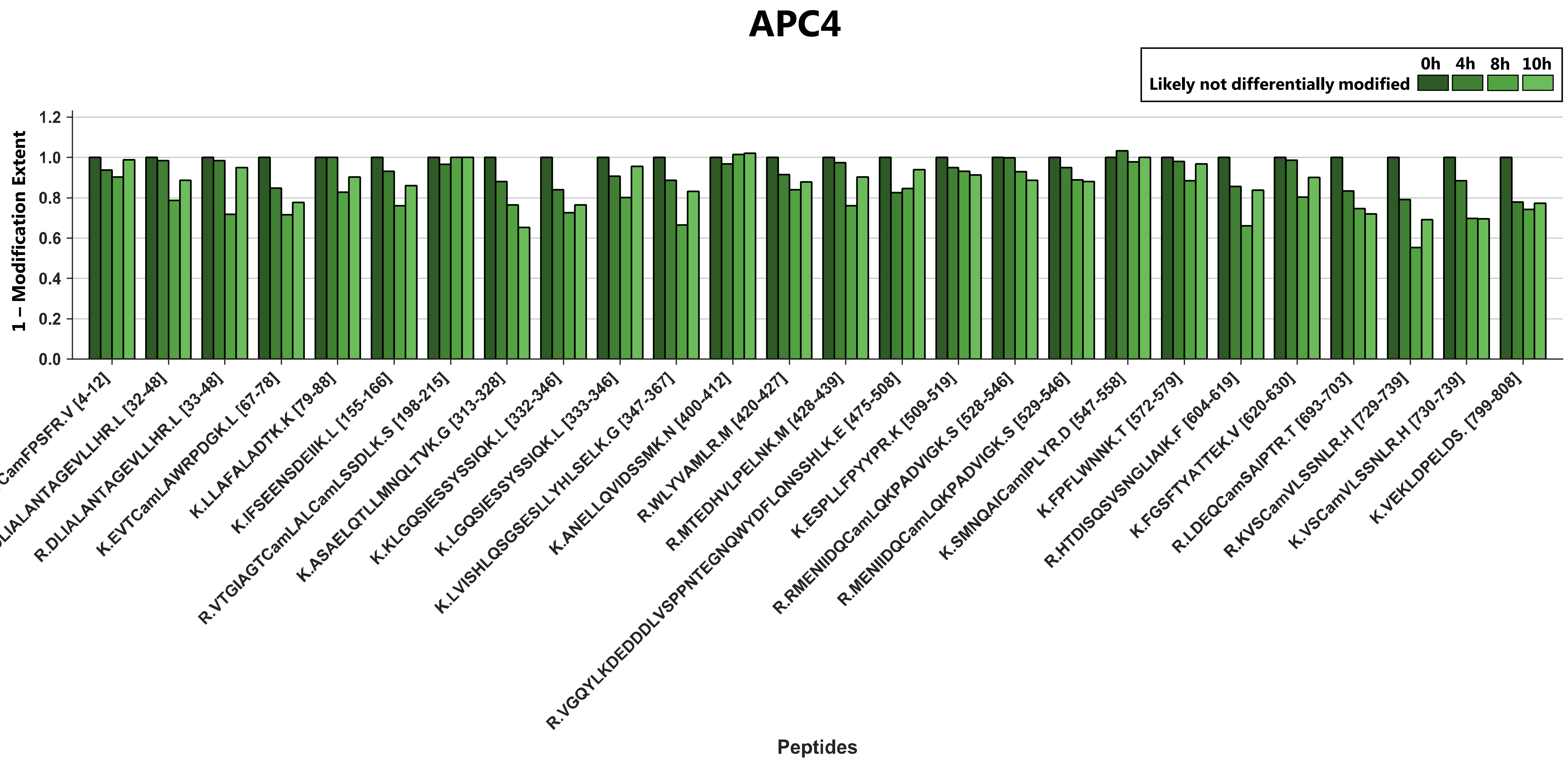
APC4
0h
4h
8h
10h
Likely not differentially modified
1 – Modification Extent
1.2
1.0
0.8
0.6
0.4
0.2
0.0
R.FPTCamFPSFR.V [4-12]
K.RDLIALANTAGEVLLHR.L [32-48]
R.DLIALANTAGEVLLHR.L [33-48]
K.EVTCamLAWRPDGK.L [67-78]
K.LLAFALADTK.K [79-88]
K.IFSEENSDEIIK.L [155-166]
R.VTGIAGTCamLALCamLSSDLK.S [198-215]
K.ASAELQTLLMNQLTVK.G [313-328]
K.KLGQSIESSYSSIQK.L [332-346]
K.LGQSIESSYSSIQK.L [333-346]
K.LVISHLQSGSESLLYHLSELK.G [347-367]
K.ANELLQVIDSSMK.N [400-412]
R.WLYVAMLR.M [420-427]
R.MTEDHVLPELNK.M [428-439]
R.VGQYLKDEDDDLVSPPNTEGNQWYDFLQNSSHLK.E [475-508]
K.ESPLLFPYYPR.K [509-519]
R.RMENIIDQCamLQKPADVIGK.S [528-546]
R.MENIIDQCamLQKPADVIGK.S [529-546]
K.SMNQAICamIPLYR.D [547-558]
K.FPFLWNNK.T [572-579]
R.HTDISQSVSNGLIAIK.F [604-619]
K.FGSFTYATTEK.V [620-630]
R.LDEQCamSAIPTR.T [693-703]
R.KVSCamVLSSNLR.H [729-739]
K.VSCamVLSSNLR.H [730-739]
K.VEKLDPELDS. [799-808]
Peptides

C

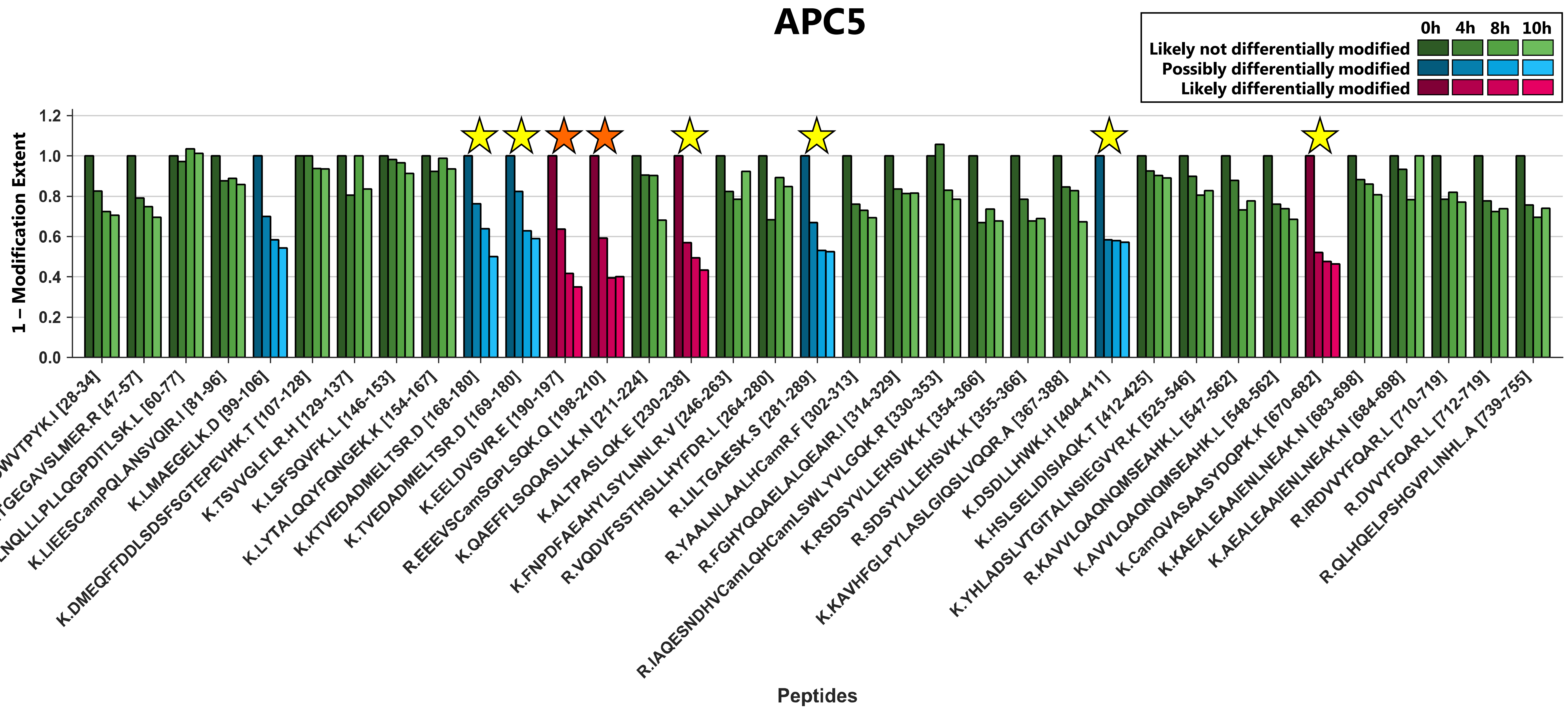
APC5
0h
4h
8h
10h
Likely not differentially modified
Possibly differentially modified
Likely differentially modified
1 – Modification Extent
1.2
1.0
0.8
0.6
0.4
0.2
0.0
K.DWVTPYK.I [28-34]
R.TGEGAVSLMER.R [47-57]
R.LNQLLLPLLQGPDITLSK.L [60-77]
K.LIEESCamPQLANSVQIR.I [81-96]
K.LMAEGELK.D [99-106]
K.DMEQFFDDLSDSFSGTEPEVHK.T [107-128]
K.TSVVGLFLR.H [129-137]
K.LSFSQVFK.L [146-153]
K.LYTALQQYFQNGEK.K [154-167]
K.KTVEDADMELTSR.D [168-180]
K.TVEDADMELTSR.D [169-180]
K.EELDVSVR.E [190-197]
R.EEEVSCamSGPLSQK.Q [198-210]
K.QAEFFLSQQASLLK.N [211-224]
K.ALTPASLQK.E [230-238]
K.FNPDFAEAHYLSYLNNLR.V [246-263]
R.VQDVFSSTHSLLHYFDR.L [264-280]
R.LILTGAESK.S [281-289]
R.YAALNLAALHCamR.F [302-313]
R.FGHYQQAELALQEAIR.I [314-329]
R.IAQESNDHVCamLQHCamLSWLYVLGQK.R [330-353]
K.RSDSYVLLEHSVK.K [354-366]
R.SDSYVLLEHSVK.K [355-366]
K.KAVHFGLPYLASLGIQSLVQQR.A [367-388]
K.DSDLLHWK.H [404-411]
K.HSLSELIDISIAQK.T [412-425]
K.YHLADSLVTGITALNSIEGVYR.K [525-546]
R.KAVVLQAQNQMSEAHK.L [547-562]
K.AVVLQAQNQMSEAHK.L [548-562]
K.CamQVASAASYDQPK.K [670-682]
K.KAEALEAAIENLNEAK.N [683-698]
K.AEALEAAIENLNEAK.N [684-698]
R.IRDVVYFQAR.L [710-719]
R.DVVYFQAR.L [712-719]
R.QLHQELPSHGVPLINHL.A [739-755]
Peptides

**D**

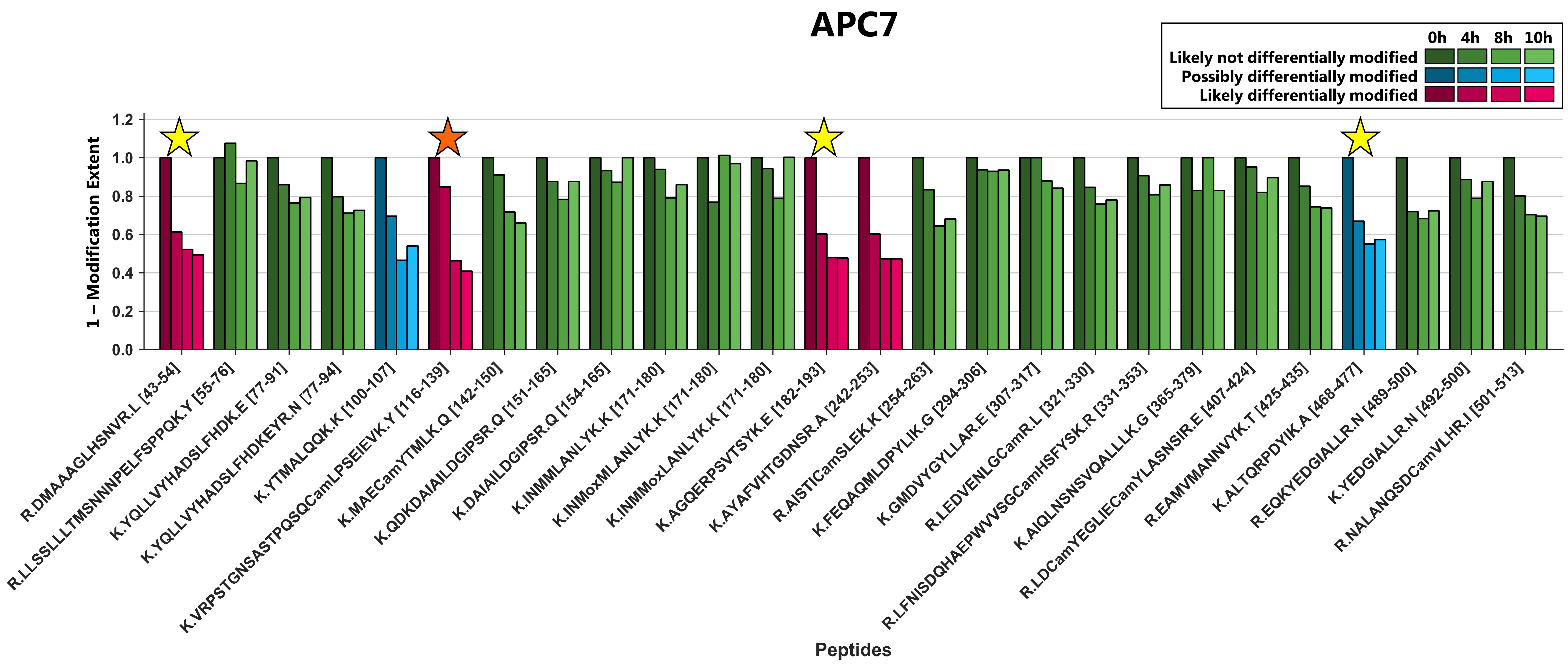

APC7
0h
4h
8h
10h
Likely not differentially modified
Possibly differentially modified
Likely differentially modified
1 – Modification Extent
1.2
1.0
0.8
0.6
0.4
0.2
0.0
R.DMAAAGLHSNVR.L [43-54]
R.LLSSLLLTMSNNNPELFSPPQK.Y [55-76]
K.YQLLVYHADSLFHDK.E [77-91]
K.YQLLVYHADSLFHDKEYR.N [77-94]
K.YTMALQQK.K [100-107]
K.VRPSTGNSASTPQSQCamLPSEIEVK.Y [116-139]
K.MAECamYTMLK.Q [142-150]
K.QDKDAIAILDGIPSR.Q [151-165]
K.DAIAILDGIPSR.Q [154-165]
K.INMMLANLYK.K [171-180]
K.INMoxMLANLYK.K [171-180]
K.INMMoxLANLYK.K [171-180]
K.AGQERPSVTSYK.E [182-193]
K.AYAFVHTGDNSR.A [242-253]
R.AISTICamSLEK.K [254-263]
K.FEQAQMLDPYLIK.G [294-306]
K.GMDVYGYLLAR.E [307-317]
R.LEDVENLGCamR.L [321-330]
R.LFNISDQHAEPWVVSGCamHSFYSK.R [331-353]
K.AIQLNSNSVQALLLK.G [365-379]
R.LDCamYEGLIECamYLASNSIR.E [407-424]
R.EAMVMANNVYK.T [425-435]
K.ALTQRPDYIK.A [468-477]
R.EQKYEDGIALLR.N [489-500]
K.YEDGIALLR.N [492-500]
R.NALANQSDCamVLHR.I [501-513]
Peptides

E

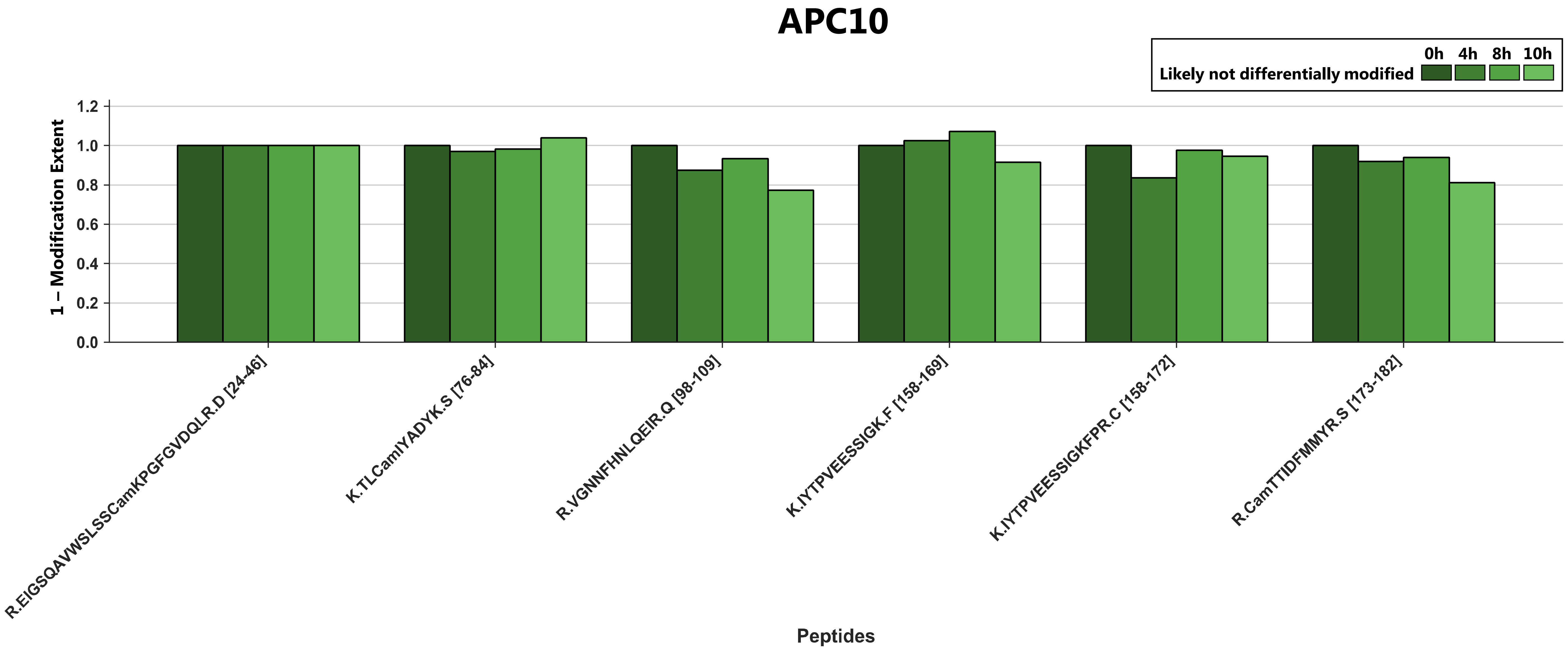

APC10
0h
4h
8h
10h
Likely not differentially modified
1.2
1.0
0.8
0.6
0.4
0.2
0.0
1 – Modification Extent
R.EIGSQAVWSLSSCamKPGFGVDQLR.D [24-46]
K.TLCamIYADYK.S [76-84]
R.VGNNFHNLQEIR.Q [98-109]
K.IYTPVEESSIGK.F [158-169]
K.IYTPVEESSIGKFPR.C [158-172]
R.CamTTIDFMMYR.S [173-182]
Peptides

F

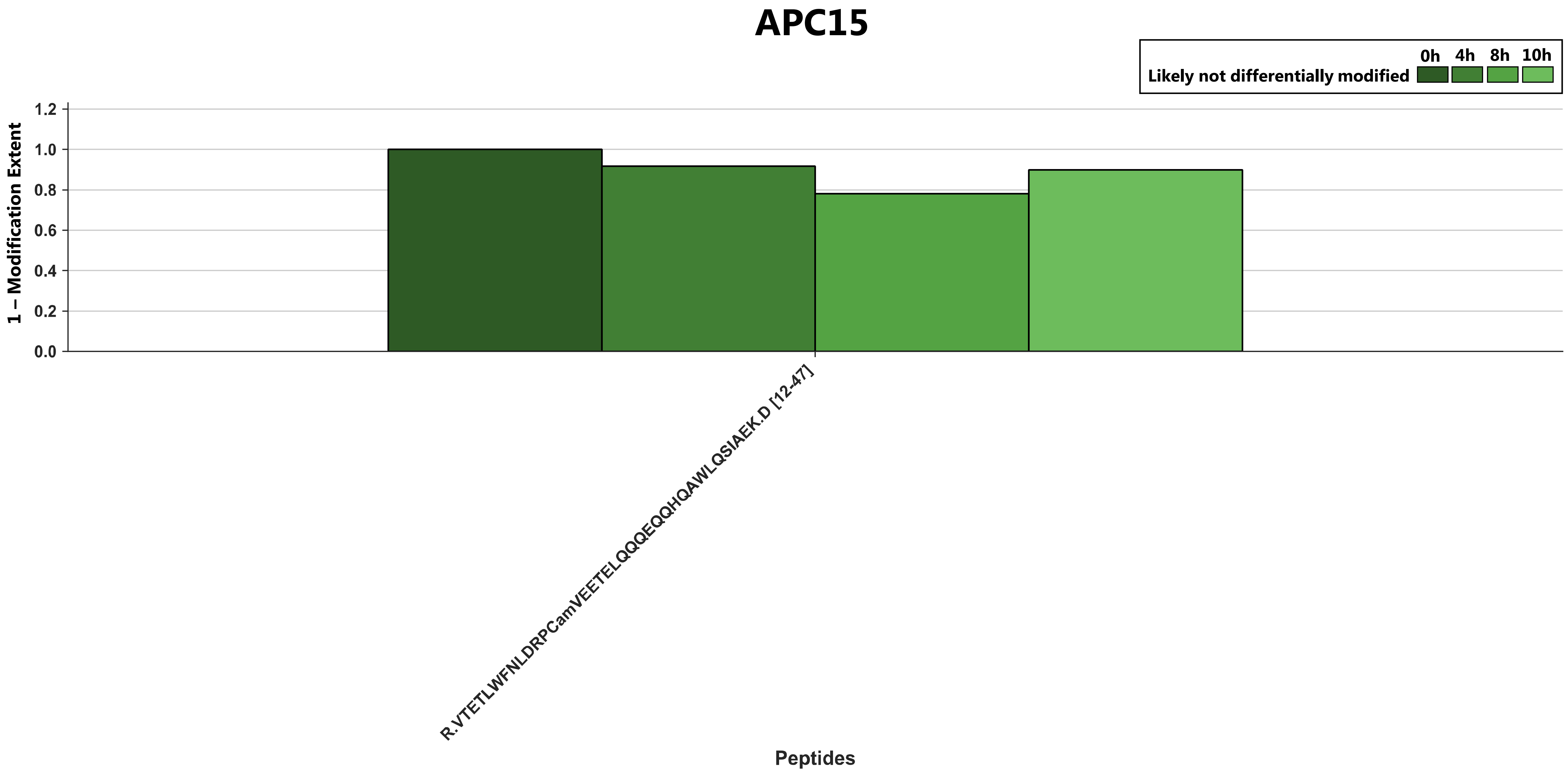

APC15
0h
4h
8h
10h
Likely not differentially modified
1 – Modification Extent
1.2
1.0
0.8
0.6
0.4
0.2
0.0
R.VTETLWFNLDRPCamVEETELQQQEQQHQAWLQSIAEK.D [12-47]
Peptides

G

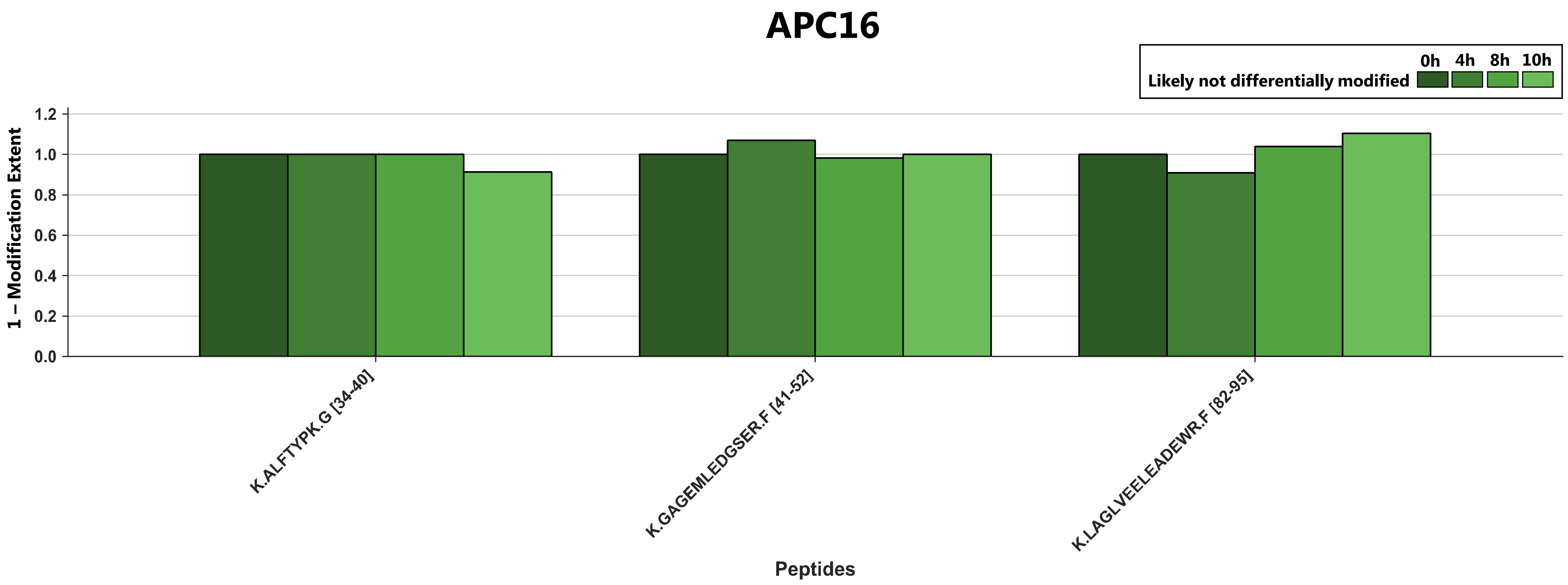

APC16
0h 4h 8h 10h
Likely not differentially modified
1 – Modification Extent
1.2
1.0
0.8
0.6
0.4
0.2
0.0
K.ALFTYPK.G [34-40]
K.GAGEMLEDGSER.F [41-52]
K.LAGLVEELEADEWR.F [82-95]
Peptides

H

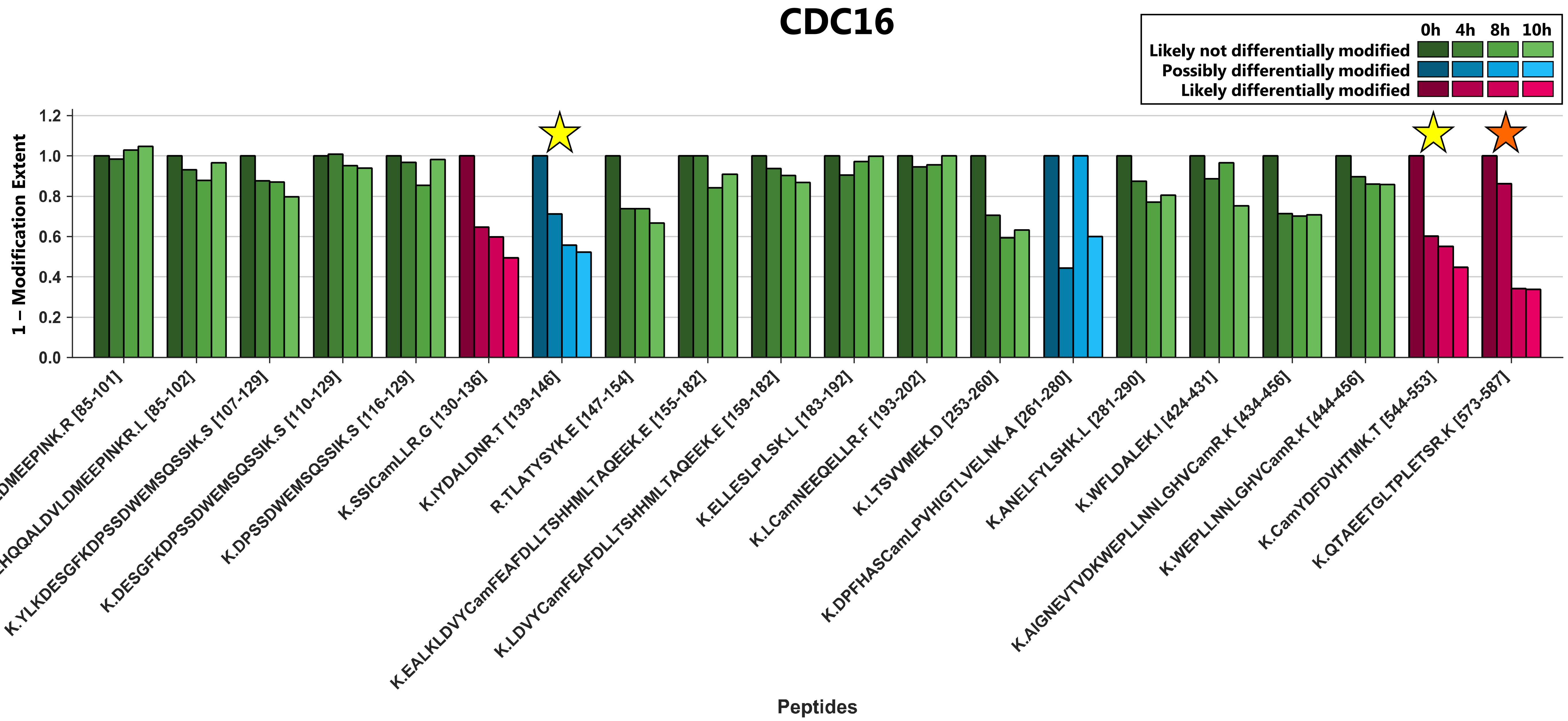

CDC16
0h 4h 8h 10h
Likely not differentially modified
Possibly differentially modified
Likely differentially modified
1 – Modification Extent
0.0
0.2
0.4
0.6
0.8
1.0
1.2
K.EHQQALDVLDMEEPINK.R [85-101]
K.EHQQALDVLDMEEPINKR.L [85-102]
K.YLKDESGFKDPSSDWEMSQSSIK.S [107-129]
K.DESGFKDPSSDWEMSQSSIK.S [110-129]
K.DPSSDWEMSQSSIK.S [116-129]
K.SSICamLLR.G [130-136]
K.IYDALDNR.T [139-146]
R.TLATYSYK.E [147-154]
K.EALKLDVYCamFEAFDLLTSHHMLTAQEEK.E [155-182]
K.LDVYCamFEAFDLLTSHHMLTAQEEK.E [159-182]
K.ELLESLPLSK.L [183-192]
K.LCamNEEQELLR.F [193-202]
K.LTSVVMEK.D [253-260]
K.DPFHASCamLPVHIGTLVELNK.A [261-280]
K.ANELFYLSHK.L [281-290]
K.WFLDALEK.I [424-431]
K.AIGNEVTVDKWEPLLNNLGHVCamR.K [434-456]
K.WEPLLNNLGHVCamR.K [444-456]
K.CamYDFDVHTMK.T [544-553]
K.QTAEETGLTPLETSR.K [573-587]
Peptides

I

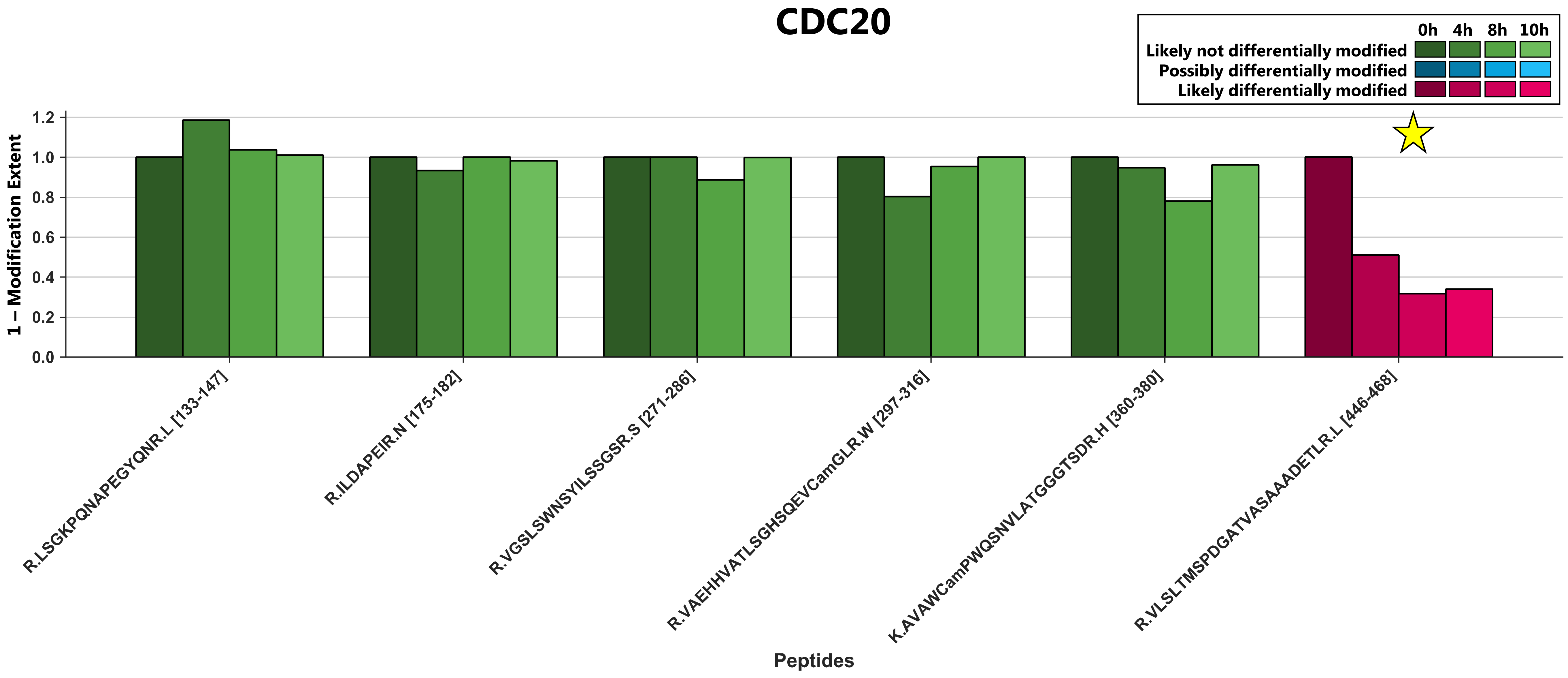

CDC20
0h
4h
8h
10h
Likely not differentially modified
Possibly differentially modified
Likely differentially modified
1 – Modification Extent
1.2
1.0
0.8
0.6
0.4
0.2
0.0
R.LSGKPQNAPEGYQNR.L [133-147]
R.ILDAPEIR.N [175-182]
R.VGSLSWNSYILSSGSR.S [271-286]
R.VAEHHVATLSGHSQEVCamGLR.W [297-316]
K.AVAWCamPWQSNVLATGGGTSDR.H [360-380]
R.VLSLTMSPDGATVASAAADETLR.L [446-468]
Peptides

J

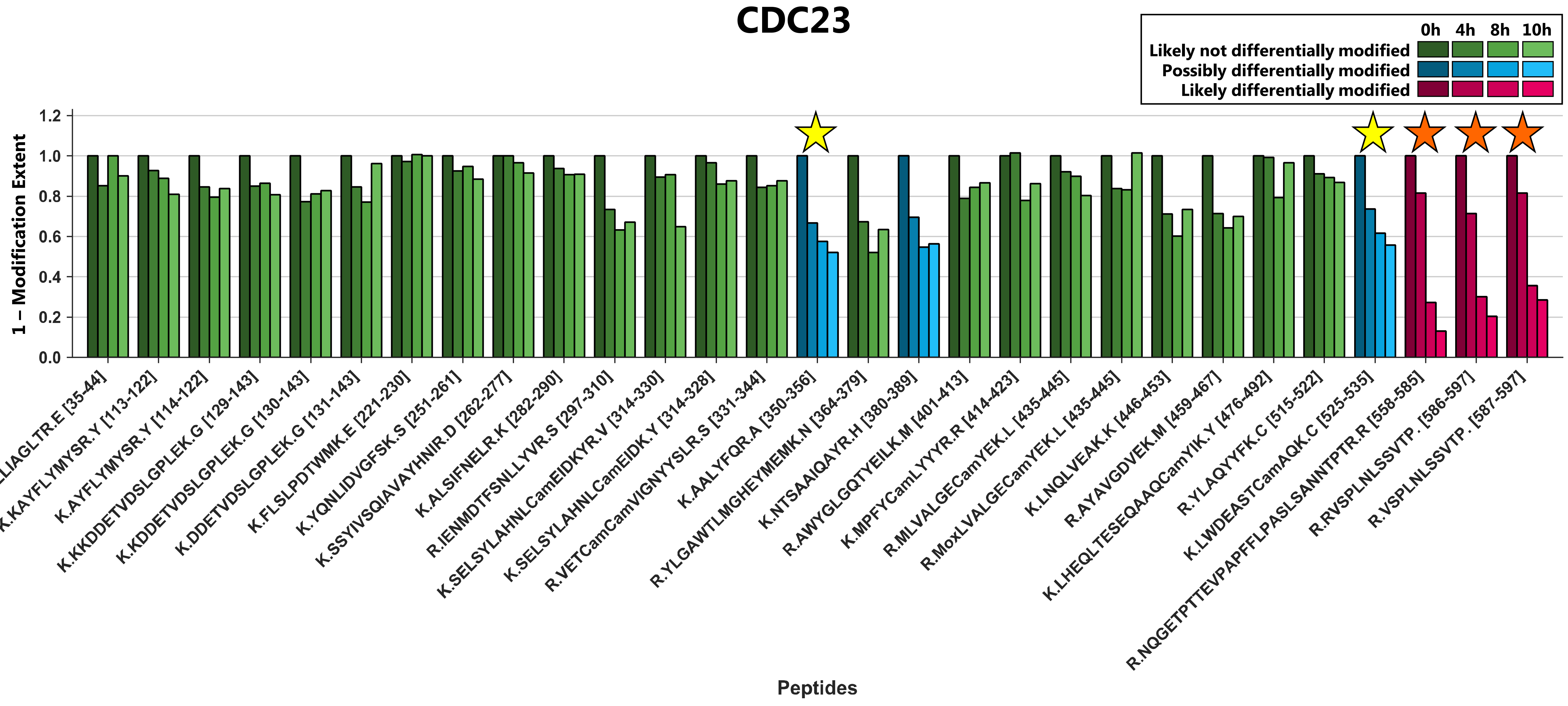

CDC23
0h
4h
8h
10h
Likely not differentially modified
Possibly differentially modified
Likely differentially modified
1 – Modification Extent
1.2
1.0
0.8
0.6
0.4
0.2
0.0
K.QLLLIAGLTR.E [35-44]
K.KAYFLYMYSR.Y [113-122]
K.AYFLYMYSR.Y [114-122]
K.KKDDETVDSLGPLEK.G [129-143]
K.KDDETVDSLGPLEK.G [130-143]
K.DDETVDSLGPLEK.G [131-143]
K.FLSLPDTWMK.E [221-230]
K.YQNLIDVGFSK.S [251-261]
K.SSYIVSQIAVAYHNIR.D [262-277]
K.ALSIFNELR.K [282-290]
R.IENMDTFSNLLYVR.S [297-310]
K.SELSYLAHNLCamEIDKYR.V [314-330]
K.SELSYLAHNLCamEIDK.Y [314-328]
R.VETCamCamVIGNYYSLR.S [331-344]
K.AALYFQR.A [350-356]
R.YLGAWTLMGHEYMEMK.N [364-379]
K.NTSAAIQAYR.H [380-389]
R.AWYGLGQTYEILK.M [401-413]
K.MPFYCamLYYYR.R [414-423]
R.MLVALGECamYEK.L [435-445]
R.MoxLVALGECamYEK.L [435-445]
K.LNQLVEAK.K [446-453]
R.AYAVGDVEK.M [459-467]
K.LHEQLTESEQAAQCamYIK.Y [476-492]
R.YLAQYYFK.C [515-522]
K.LWDEASTCamAQK.C [525-535]
R.NQGETPTTEVPAPFFLPASLSANNTPTR.R [558-585]
R.RVSPLNLSSVTP . [586-597]
R.VSPLNLSSVTP . [587-597]
Peptides

K

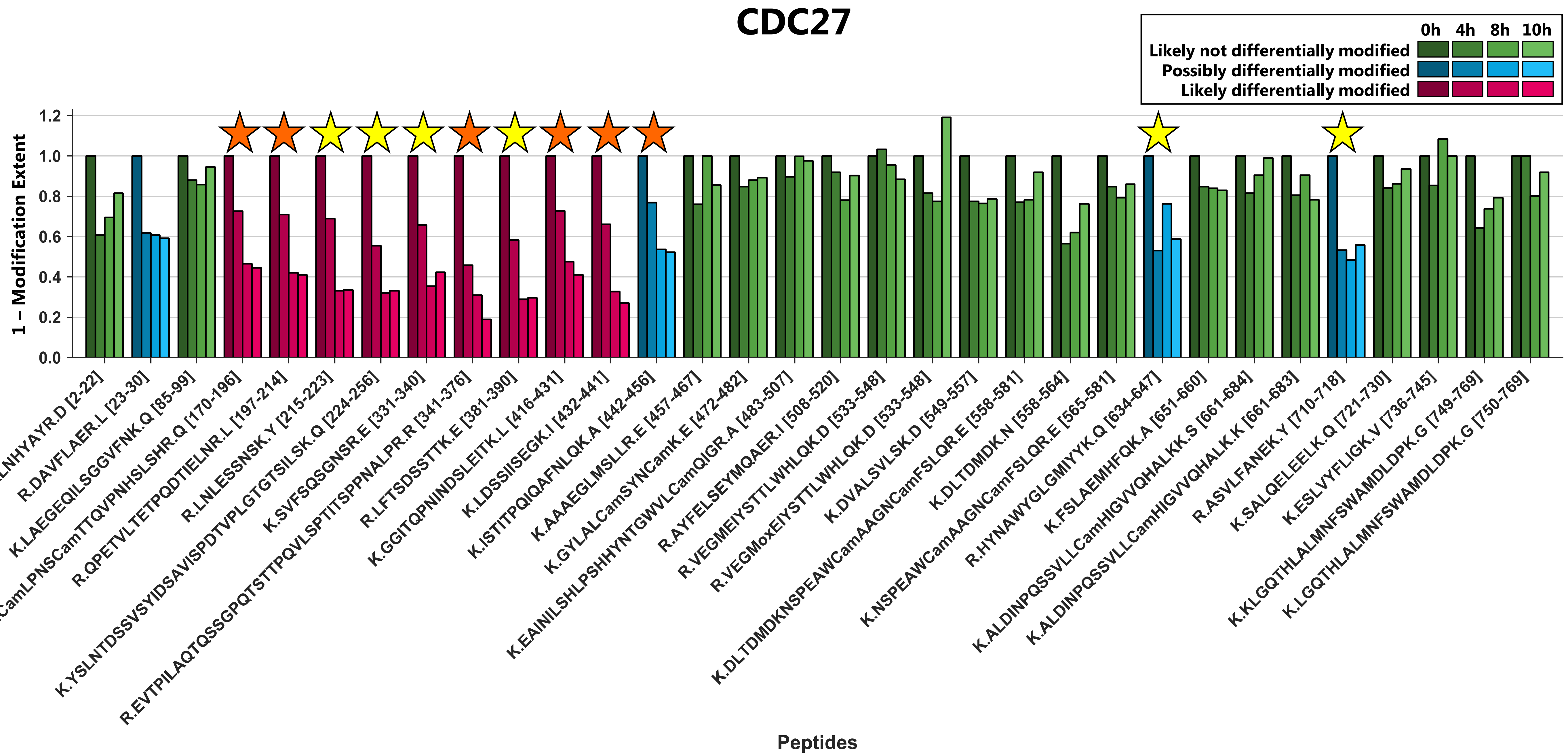
CDC27
0h 4h 8h 10h
Likely not differentially modified
Possibly differentially modified
Likely differentially modified
1 – Modification Extent
1.2
1.0
0.8
0.6
0.4
0.2
0.0
M.TVLQEPVQAAIWQALNHYAYR.D [2-22]
R.DAVFLAER.L [23-30]
K.LAEGEQILSGGVFNK.Q [85-99]
K.FTSLQNFSNCamLPNSCamTTQVPNHSLSHR.Q [170-196]
R.QPETVLTETPQDTIELNR.L [197-214]
R.LNLESSNSK.Y [215-223]
K.YSLNTDSSVSYIDSAVISPDTVPLGTGTSILSK.Q [224-256]
K.SVFSQSGNSR.E [331-340]
R.EVTPILAQTQSSGPQTSTTPQVLSPTITSPPNALPR.R [341-376]
R.LFTSDSSTTK.E [381-390]
K.GGITQPNINDSLEITK.L [416-431]
K.LDSSIISEGK.I [432-441]
K.ISTITPQIQAFNLQK.A [442-456]
K.AAAEGLMSLLR.E [457-467]
K.GYLALCamSYNCamK.E [472-482]
K.EAINILSHLPSHHYNTGWVLCamQIGR.A [483-507]
R.AYFELSEYMQAER.I [508-520]
R.VEGMEIYSTTLWHLQK.D [533-548]
R.VEGMoxEIYSTTLWHLQK.D [533-548]
K.DVALSVLSK.D [549-557]
K.DLTDMDKNSPEAWCamAAGNCamFSLQR.E [558-581]
K.DLTDMDK.N [558-564]
K.NSPEAWCamAAGNCamFSLQR.E [565-581]
R.HYNAWYGLGMIYYK.Q [634-647]
K.FSLAEMHFQK.A [651-660]
K.ALDINPQSSVLLCamHIGVVQHALK.S [661-684]
K.ALDINPQSSVLLCamHIGVVQHALK.K [661-683]
R.ASVLFANEK.Y [710-718]
K.SALQELEELK.Q [721-730]
K.ESLVYFLIGK.V [736-745]
K.KLGQTHLALMNFSWAMDLDPK.G [749-769]
K.LGQTHLALMNFSWAMDLDPK.G [750-769]
Peptides
